# Soaring styles of extinct giant birds and pterosaurs

**DOI:** 10.1101/2020.10.31.354605

**Authors:** Yusuke Goto, Ken Yoda, Henri Weimerskirch, Katsufumi Sato

## Abstract

The largest extinct volant birds (*Pelagornis sandersi* and *Argentavis magnificens*) and pterosaurs (*Pteranodon* and *Quetzalcoatlus*) are thought to have used wind-dependent soaring flight, similar to modern large birds. There are two types of soaring: thermal soaring, used by condors and frigatebirds, which involves the use of updrafts to ascend and then glide horizontally over the land or the sea; and dynamic soaring, used by albatrosses, which involves the use of wind speed differences with height above the sea surface. Previous studies have suggested that *Pelagornis sandersi* used dynamic soaring, while *Argenthavis magnificens, Pteranodon*, and *Quetzalcoatlus* used thermal soaring. However, the performance and wind speed requirements of dynamic and thermal soaring for these species have not yet been quantified comprehensively. We quantified these values using aerodynamic models and compared them with that of extant birds. For dynamic soaring, we quantified maximum flight speeds and maximum upwind flight speeds. For thermal soaring, we quantified the animal’s sinking speed circling at a given radius and how far it could glide losing a given height. Our results confirmed those from previous studies that *Pteranodon* and *Argentavis magnificens* used thermal soaring. Conversely, the results for *Pelagornis sandersi* and *Quetzalcoatlus* were contrary to those from previous studies. *Pelagornis sandersi* used thermal soaring, and *Quetzalcoatlus* had a poor ability both in dynamic and thermal soaring. Our results demonstrate the need for comprehensive assessments of performance and required wind conditions when estimating soaring styles of extinct flying species.

## Introduction

Flying animals have evolved a wide range of body sizes. Among them, there have been incredibly large species of birds and pterosaurs (Fig. 1). *Pelagornis sandersi* and *Argentavis magnificens* are the largest extinct volant birds. Their estimated wingspans reached 6–7 m (*1*–*4*), twice as large as that of the wandering albatross, the extant bird with the longest wingspan (Table 1). Several large species of pterosaurs appeared in the Cretaceous period. *Pteranodon,* presumably the most famous pterosaur, is estimated to have had a wingspan of 6 m (Table 1) (*5*). The azhdarchids are one of the most successful Cretaceous pterosaur groups and include several large species with wingspans of approximately 10 m (Table 1) (*6*–*9*). *Quetzalcoatlus northorpi*, an azhdarchid species, is regarded as one of the largest flying animals in history.

**Fig. 1.**
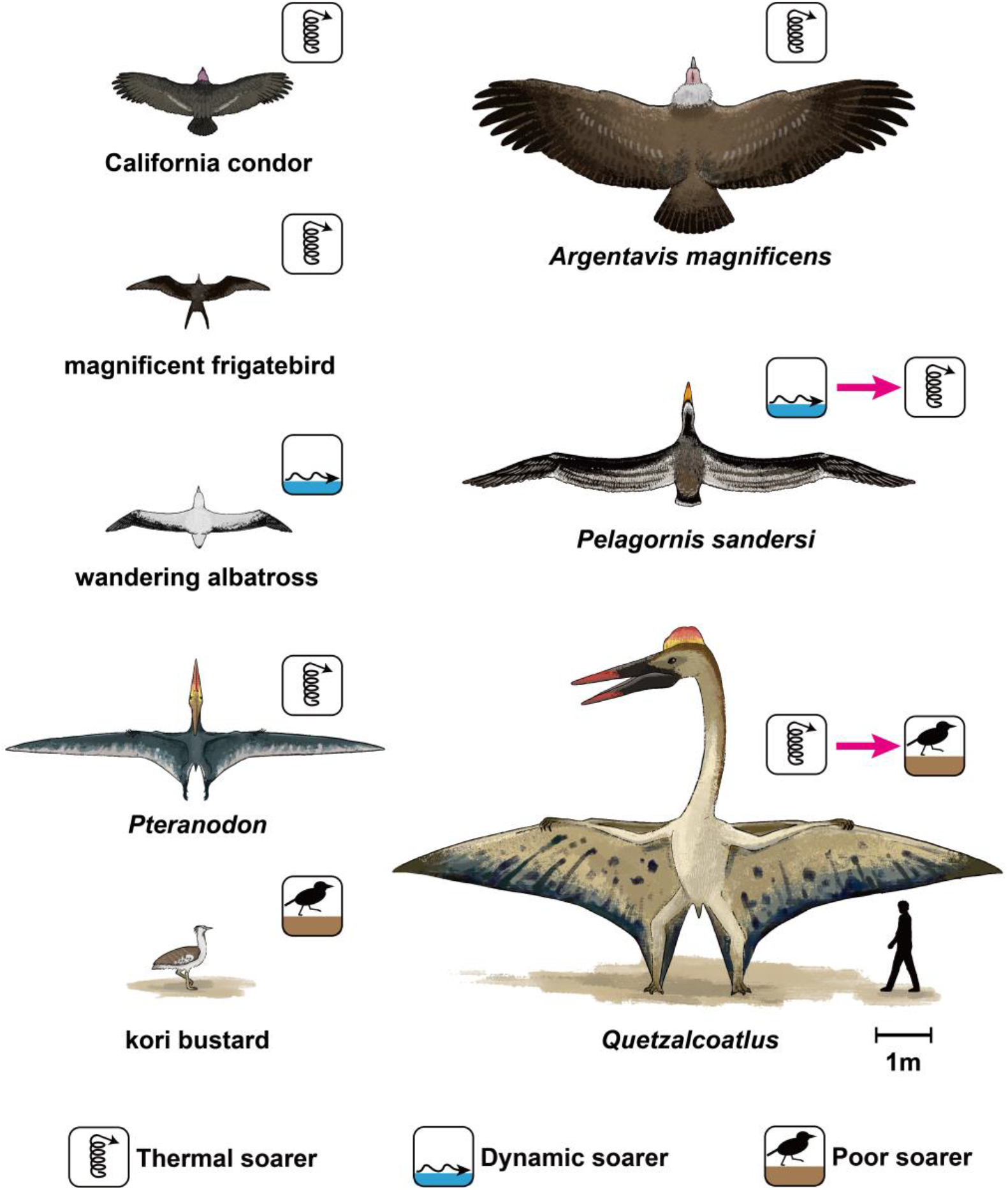
A size comparison and soaring styles of large flying animals. A size comparison and soaring styles of extinct giant birds (*Pelagornis sandersi* and *Argentavis magnificens*), pterosaurs (*Pteranodon* and *Quetzalcoatlus*), the largest extant dynamic soaring bird (wandering albatross), the largest extant thermal soaring terrestrial bird (California condor), a large extant thermal soaring seabird (magnificent frigatebird), and the heaviest extant volant bird (kori bustard). The icons indicate dynamic soarer, thermal soarer, and poor soarer and summarize the main results of this study. The red arrows indicate the transition from a previous expectation or hypothesis to the knowledge updated in this study.

**Table 1.** Morphological values of examined species. *For the wandering albatross and the black-browed albatross, we used the averages calculated from the morphological values of males and females in the cited references (*63*, *64*). For the kori bustard, we used the morphology data available in the *Flight* program (*23*).

| Species |  | Mass<br>(kg) | Wingspan<br>(m) | Wing area<br>(m<sup>2</sup>) | Aspect<br>ratio | Wing loading<br>[N/m<sup>2</sup>] | Ref |
| --- | --- | --- | --- | --- | --- | --- | --- |
| Extinct | *Pelagornis sandersi* | 21.8,<br>and<br>40.1 | 6.06, 6.13,<br>6.40 and<br>7.38 | 2.45<br>~<br>4.19 | 13.0,<br>14.0,<br>and<br>15.0 | 51.0 ~ 87.4<br>(with 21.8 kg Mass)<br>93.9 ~ 161<br>(with 40.1 kg Mass) | (1) |
|  | *Argentavis magnificens* | 70.0 | 7.00 | 8.11 | 6.04 | 84.7 | (4) |
|  | *Pteranodon* | 36.7 | 5.96 | 1.99 | 17.9 | 181 | (5) |
|  | *Quetzalcoatlus* | 259 | 9.64 | 11.4 | 8.18 | 224 | (5) |
| Dynamic soaring | wandering albatross<br>(*Diomedea exulans*) | 8.64 | 3.05 | 0.606 | 15.4 | 140 | (63)* |
|  | black-browed albatross<br>(*Thalassarche melanophris*) | 3.55 | 2.25 | 0.376 | 13.4 | 87.5 | (64)* |
|  | white-chinned petrel<br>(*Procellaria aequinoctialis*) | 1.37 | 1.40 | 0.169 | 11.6 | 79.5 | (65) |
| Thermal soaring | magnificent frigatebird<br>(*Fregata magnificens*) | 1.52 | 2.29 | 0.408 | 12.8 | 36.5 | (24) |
|  | California condor<br>(*Gymnogyps californianus*) | 9.50 | 2.74 | 1.32 | 5.70 | 70.6 | (4) |
|  | brown pelican<br>(*Pelecanus occidentalis*) | 2.65 | 2.10 | 0.450 | 9.80 | 57.8 | (24) |
|  | black vulture<br>(*Coragyps atratus*) | 1.82 | 1.38 | 0.327 | 5.82 | 54.6 | (24) |
|  | white stork<br>(*Ciconia ciconia*) | 3.40 | 2.18 | 0.540 | 7.42 | 61.8 | (4) |
| Non-Soaring | kori bustard<br>(*Ardeotis kori*) | 11.9 | 2.47 | 1.06 | 5.76 | 110 | (23) |
| Motor<br>glider | Schleicher ASK-14 | 340 | 14.3 | 12.6 | 16.2 | 265 | (33) |

How and how well these giant animals were able to fly has fascinated researchers across disciplines for over a century (*10*). This is because the question is not only interesting as a biophysical question in its own right, but also because it contributes to unraveling a wide range of issues such as the lifestyle of these species, their role in paleoecosystems, and the drivers of morphological evolution, diversification, and extinction of giant species over geological time (*11*–*15*). Their huge size must have significantly affected their flight because, with increasing size, the power required to fly increases faster than the power muscles can produce via the flapping of wings (*16*). Hence, this physical constraint has resulted in two heated arguments about the flight of extinct giants. The first is about whether and how they were able to take off (*4*, *17*–*19*). The present study focuses on the second argument. Due to the high costs of flapping that stems from their large body size, large extant birds prefer to fly utilizing wind energy or convection, that is, they prefer to soar (*17*, *20*). Hence, it is presumed that extinct large animals also employed soaring flight as their primary mode of transportation (*1*, *4*, *13*). The second argument is about what kind of soaring flight style they employed (*1*, *4*, *13*, *21*, *22*).

There are two main soaring flight styles among extant birds: dynamic soaring and thermal soaring (*23*). In dynamic soaring, birds extract flight energy from wind shear—the vertical gradient in horizontal wind speed over the ocean (Fig. 2A). Extant seabirds (e.g., albatrosses, shearwaters, and petrels) employ this soaring style and can routinely travel hundreds of kilometers per day over the sea. In thermal soaring, birds first fly circling in warm rising-air columns (thermals). They climb to a substantial height and then glide off in the desired direction while losing their height (Fig. 2C–E). By repeating this up-down process, birds travel over vast distances. Various terrestrial bird species (e.g., vultures, eagles, and storks) and seabirds (e.g., frigatebirds and pelicans) employ thermal soaring (*24*).

**Fig. 2.**
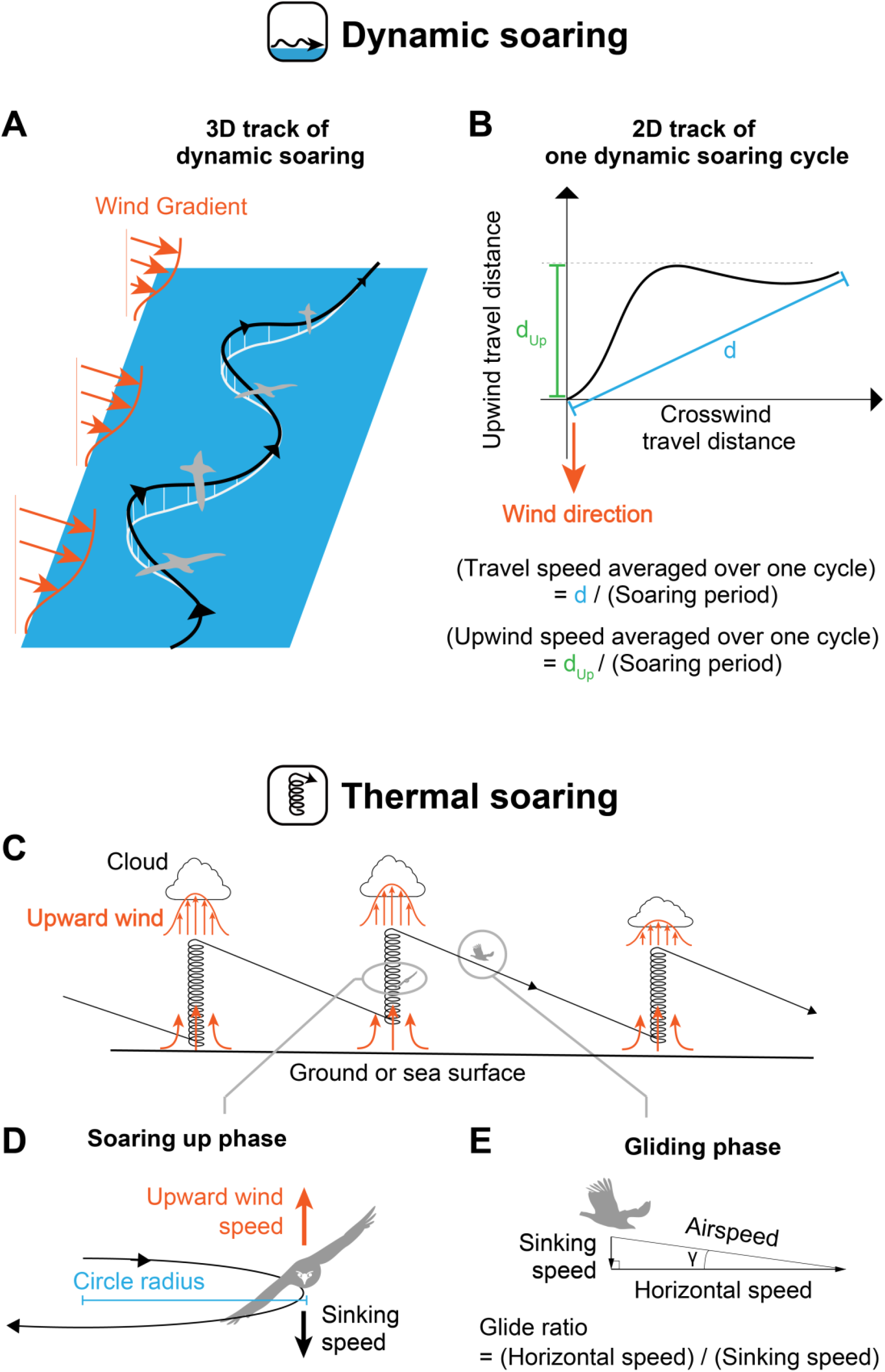
Schematics of dynamic soaring and thermal soaring. (A) Example of a 3D track of dynamic soaring. Dynamic soaring species repeat an up and down process with a shallow S-shaped trajectory at the sea surface. By utilizing wind gradients, a species can fly without flapping. (B) Example of a 2D dynamic soaring trajectory of one soaring cycle. The travel speed averaged over one cycle is defined as the travel distance in one cycle (d) divided by the soaring period, and the upwind speed averaged over one cycle is defined as the upwind travel distance in one cycle (dUp) divided by the soaring period. (C) Schematic of a thermal soaring cycle. (D) In the soaring up phase, a species soars in a steady circle. When there is upward wind that is greater than a species’ sinking speed, the species can ascend in the thermal. The upward wind is stronger in the center of a thermal; therefore, achieving a small circle radius is advantageous for thermal soaring. (E) In the gliding phase a species glides in a straight line. The rate of horizontal speed to the sinking speed is equal to the rate of horizontal distance traveled to the height lost.

Previous studies estimated that *Pelagornis sandersi* was a dynamic soarer, and *Argentavis magnificens*, *Pteranodon*, and *Quetzalcoatlus* were thermal soarers (*1*, *4*, *13*, *22*). See **Materials and Methods** *“Quantification of soaring styles in previous studies”* for details about previous studies on this topic. To estimate the potential soaring styles of these extinct animals, it is essential to quantify their soaring performance, e.g., potential speed and efficiency of soaring, as well as the required wind speed to sustain soaring flight. Valuable indicators of dynamic soaring performance are the maximum travel speed and the maximum upwind speed averaged over one dynamic soaring cycle (Fig. 2B) (*25*–*27*). Additionally, it is essential to evaluate the minimum horizontal wind speed required for sustainable dynamic soaring (*28*). Thermal soaring performance is well quantified by two indicators: the glide ratio, i.e., the ratio of the distance the animal traverses to the height the bird loses to cover that distance in the gliding phase (Fig. 2E), and the sinking speed of the animal circling in a given radius during the upward soaring phase (Fig. 2D). This sinking speed during circling corresponds to the upward wind speed required to ascend in a thermal. Because thermals have a stronger updraft in the center (Fig. 2C), the animal needs to achieve not only low sinking speed but also a narrow circle radius to efficiently ascend using a thermal.

However, the soaring performances and required wind conditions have not been comprehensively evaluated for *Pelagornis sandersi*, *Argentavis magnificens*, *Pteranodon*, or *Quetzalcoatlus* (summarized in Table 2). Three knowledge gaps are highlighted in Table 2: (1) the dynamic soaring performance and the required minimum wind speed have rarely been evaluated; (2) the thermal soaring performance in the soaring up phase and the required updraft wind speed have not been evaluated for *Pelagornis sandersi*, *Pteranodon*, and *Quetzalcoatlu*s; and (3) despite recent studies showing that the body masses of *Pteranodon* and *Quetzalcoatlus* were approximately three times heavier than previously expected (*5*, *17*, *29*), and that pterosaurs’ wings had a higher profile drag than that of birds (*22*), the soaring performances of these new heavy body masses and higher drags have rarely been evaluated.

**Table 2.**
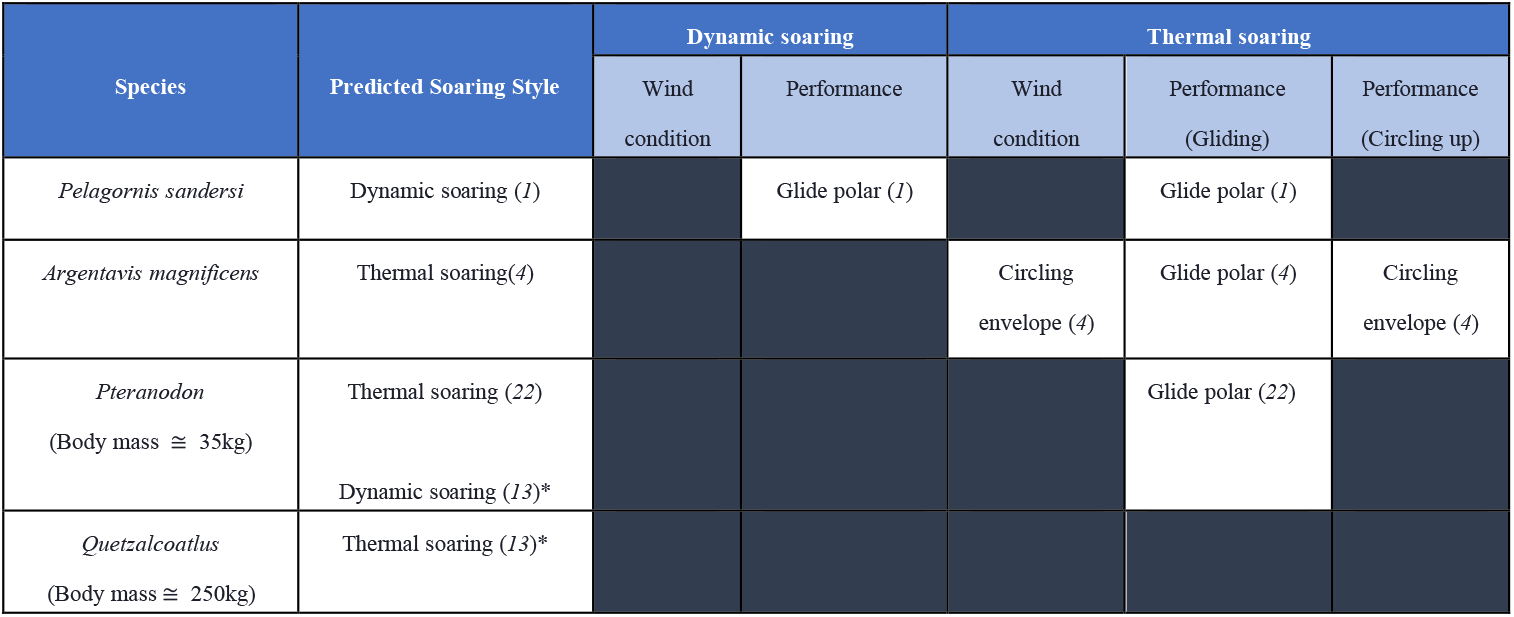
Summary of previous studies. Previous studies that quantified the soaring performances and required wind conditions of *Pelagornis sandersi*, *Argentavis magnificens*, *Pteranodon*, and *Quetzalcoatlus* with recent heavy body mass estimates. *A principal component analysis (PCA) using three morphological information (logarithm of each of the animal’s weight, wing area, and wingspan) has been performed in the previous studies for birds (*37*), bats (*36*) and pterosaurs (*5*) respectively. In the previous study (*13*), these second and third principal components were compared among birds, bats and pterosaurs to estimate the soaring styles of pterosaurs.

In this study, we aimed to address these knowledge gaps and identify the potential soaring styles of these extinct giant birds and pterosaurs. To this end, we used physical models and recent morphology estimates to quantify the performance and wind conditions required for dynamic and thermal soaring in these animals and compared them with those of extant soaring birds. See **Materials and Methods** *“Models”* for details about the employed models and parameter values.

## Results

### Dynamic soaring

We quantified the dynamic soaring performance (Fig. 2 B) and required wind speeds using a physical model and a numerical optimization method. This method has been developed in the engineering field and provides a framework to quantify dynamic soaring performances and required wind conditions for gliders and birds (*25*, *28*, *30*, *31*). However, despite its effectiveness, the only animal to which this technique has been applied is the wandering albatross (*28*, *31*); it has never been applied to extinct giant flyers. We applied this framework to the four giant extinct species and three extant dynamic soaring bird species with various sizes ranging from 1 to 9 kg [i.e., the white-chinned petrel (*Procellaria aequinoctialis*), the black-browed albatross (*Thalassarche melanophris*), and the wandering albatross (*Diomedea exulans*)]. As the exact shape of the wind gradient remains poorly understood, we conducted the calculation under seven different wind conditions (Fig. 3A–C) (*28*, *31*). In addition, we added an important modification to the previous models: the animal’s wings do not touch the sea surface during their flight (Fig. 3D; and see Eq. 16 and its description in **Materials and Methods** for details).

**Fig. 3.**
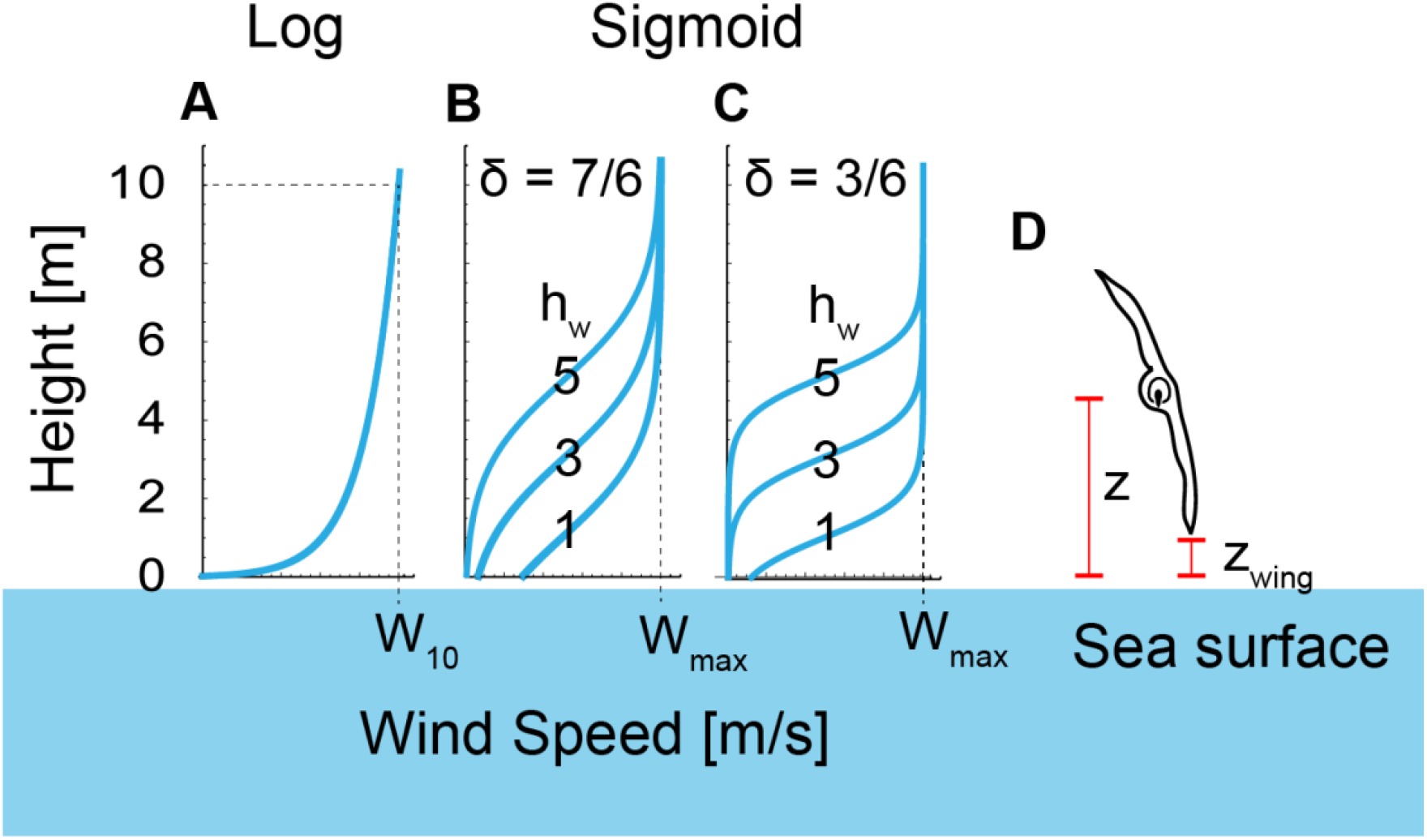
Wind shear models explored in this study. (A) Logarithmic wind gradient model. The wind speed at height 10 m was defined as *W*_10_. (B) Sigmoidal wind shear model with a wind shear thickness of 7 m (δ = 7/6) and a shear height (*h*_w_) of 1, 3, or 5 m. (C) Sigmoidal wind shear model with a wind shear thickness of 3 m (δ = 3/6) and a shear height of 1, 3, or 5 m. The maximum wind speed of the sigmoidal model is represented as W_max_. (D) Schematic of a soaring bird. Its height from the sea surface is represented as z and the height of the wingtip is represented as z_wing_. We constrained the models so that the wing tip did not touch the sea surface, i.e., *z*_wing_ ≥ 0.

Our computation results indicate that *Argentavis magnificens* and *Quetzalcoatlus* could not have employed dynamic soaring (Fig. 4 A–C). Both species showed lower dynamic soaring performances and higher required wind speeds for dynamic soaring than the extant dynamic soaring species under all wind conditions tested in this study. The dynamic soaring performances and required wind speeds of *Pelagornis sandersi* and *Pteranodon* varied substantially with the assumed morphology and shape of the wind gradient, especially the height where the steep wind speed change occurs (Fig. 4 A–C).

**Fig. 4.**
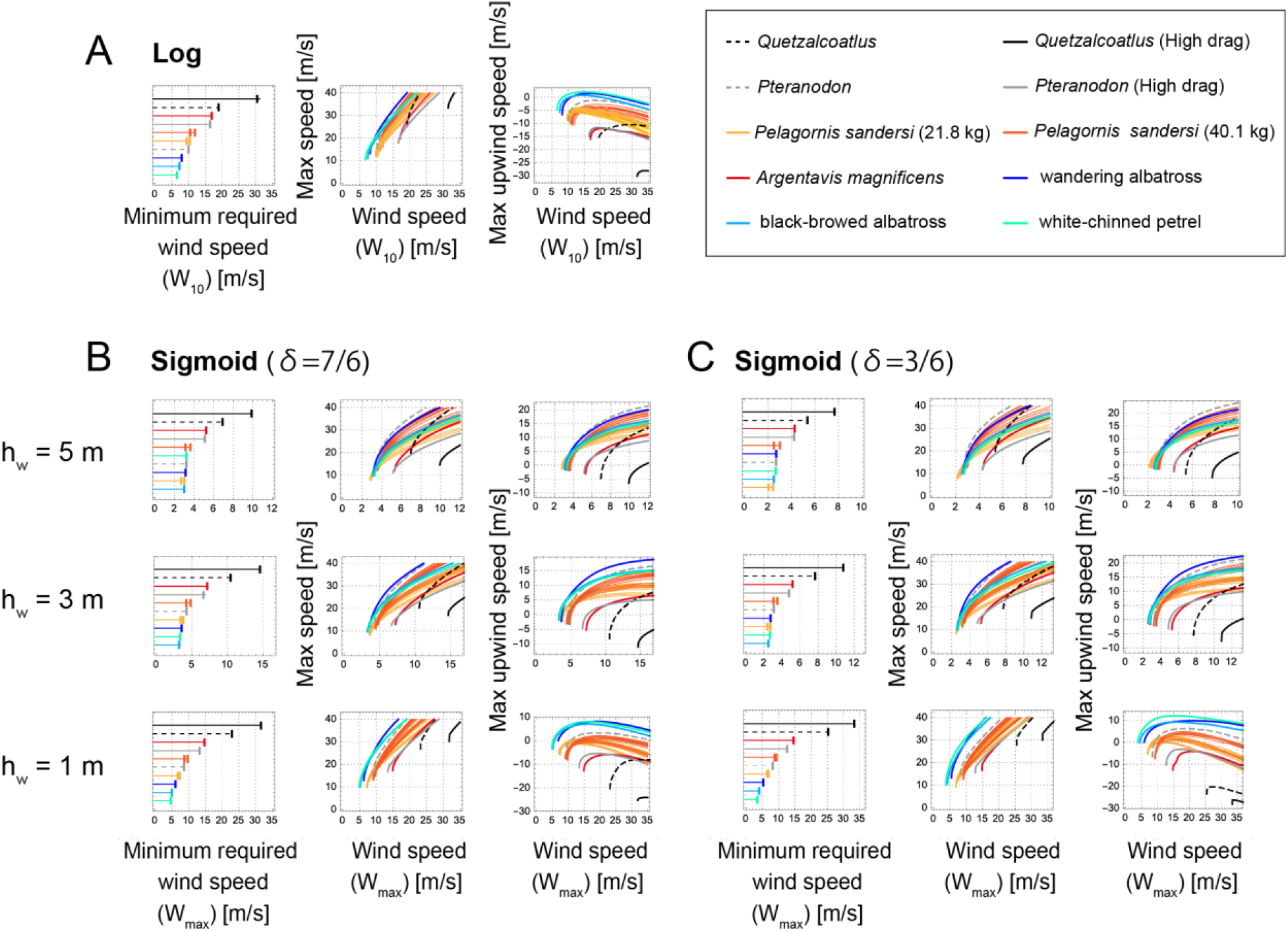
Required minimum wind speeds and dynamic soaring performances of extinct and extant animals. (A) Results of the logarithmic wind model. (B) Results of the sigmoidal wind model with a wind shear thickness of 7 m (δ = 7/6) and wind shear height (*h*_w_) of 1, 3, or 5 m. (C) Results of the Sigmoidal wind model with a wind shear thickness of 3 m (δ = 3/6) and a wind shear height (*h*_w_) of 1, 3, or 5 m. The first column shows the minimum required wind speed for sustainable dynamic soaring. The second column shows the maximum travel speed averaged over one soaring cycle, in response to wind speed. The third column shows the maximum upwind speed averaged over one soaring cycle, in response to wind speed.

When *Pteranodon* was analyzed with a high-profile drag coefficient based on a wind tunnel experiment (*22*), it showed poor performance and required strong winds compared with extant species, which suggested that *Pteranodon* did not employ dynamic soaring. When *Pteranodon* was analyzed with the same low-profile drag coefficient of birds, its performance was better than that of extant birds when the wind change was located far (upwards) from the sea surface (sigmoidal wind condition with *h*_w_ = 5 and 3). When the wind speed change was located close to the sea surface (logarithmic model and sigmoidal wind condition with *h*_w_ = 1), *Pteranodon* showed a poor flight performance. It required a stronger wind speed than the extant dynamic soaring species, except for under sigmoidal wind conditions with *h*_w_ = 5. Hence, *Pteranodon* could not employ dynamic soaring when a high-profile drag was assumed. Conversely, when a low-profile drag was assumed, *Pteranodon* was capable of dynamic soaring but required strong wind conditions.

For *Pelagornis sandersi*, the results were highly dependent on body mass. With heavy body mass estimates (40.1 kg), *Pelagornis sandersi* required higher wind speeds than extant dynamic soaring species, irrespective of the wind conditions. The performance was superior to extant species for some morphology estimates when the shear height was far from the sea surface (sigmoidal wind condition with *h*_w_ = 5 and 3 m), but inferior when the wind speed change was located close to the sea surface (logarithmic model and sigmoidal wind condition with *h*_w_ = 1). When lower body mass estimates were used (21.8 kg), *Pelagornis sandersi* required lower wind speeds, but its performance was distinctively lower than that of extant species. Hence, *Pelagornis sandersi* required harsh wind conditions for dynamic soaring when a 40.1 kg body mass was assumed, and it was poor at dynamic soaring when a 21.8 kg body mass was assumed.

The performances of all species varied with the value of *h*_w_, and the variation was especially distinct for large species in contrast to that of white-chinned petrels (Fig. 4 B and C). This variation was due to wingtip boundary conditions (Fig. 3D and Eq. [16]). Animals can attain more energy when passing through large wind speed gradients, but when large gradient changes are close to the sea level, large animals are unable to use the wind speed gradient efficiently because their wings limit the altitude available to them. Although the long, thin wings that reduce drag in extant dynamic soaring birds are suited for dynamic soaring (*23*, *32*), our detailed dynamic models have shown that excessively long wings can also inhibit efficient dynamic soaring.

### Thermal soaring

The thermal soaring performances and the required upward wind speeds (Fig. 2D and E) were quantified using the established framework, i.e., glide polars and circling envelopes. A glide polar is a graphical plot of the sinking speed versus horizontal speed when a bird glides in a straight line (*23*). We can determine the maximum glide ratio and the associated travel speed of flyers by identifying the line that passes through the origin and tangents of the glide polar plot. The inverse of the line slope and the speed at the tangent point correspond to the maximum glide ratio and the associated horizontal speed, respectively. The circling envelope is a graphical plot of the sinking speed (i.e., equivalent to the required upward wind speed for ascent in a thermal) versus the radius of a turn when a bird glides in a steady circle (*23*, *24*). We quantified the thermal soaring performances and the required upward wind speeds for the four extinct species, five extant thermal soaring species [the magnificent frigatebird (*Fregata magnificens*), the black vulture (*Coragyps atratus*), the brown pelican (*Pelecanus occidentalis*), the white stork (*Ciconia ciconia*), and the California condor (*Gymnogyps californianus*)], and the kori bustard (*Ardeotis kori*), the heaviest extant volant bird that does not soar. The performances reported in a previous study of the *Schleicher* ASK-14, a motor glider with 14 m wingspan similar to that of *Quetzalcoatlus*, are also presented for comparison (*33*).

Extinct species, expect for *Quetzalcoatlus*, showed high gliding performances with maximum glide ratios ranging from 11 to 22 (Fig. 5A), which are comparative to those of extant species (from 11 to 18); nevertheless, *Quetzalcoatlus* had the lowest soaring efficiency (i.e., 8) of the species when a high drag coefficient was assumed.

**Fig. 5.**
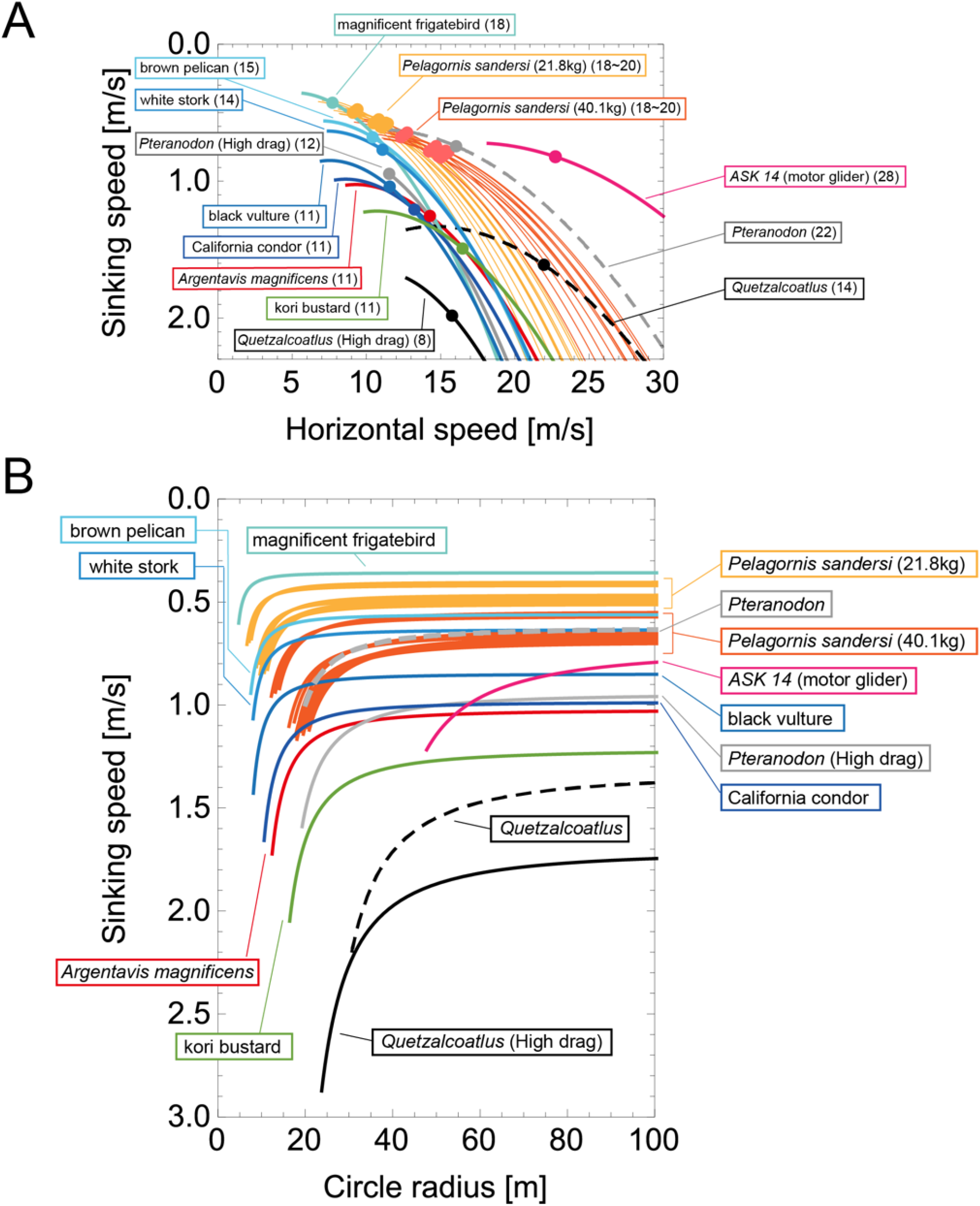
Glide polars (A) and circling envelopes (B) of extinct species, extant thermal soaring species, and the kori bustard, the heaviest rarely flying bird. The dashed line of *Pteranodon* and *Quetzalcoatlus* represents the result of a low-profile drag coefficient (CD_pro_ = 0.014), similar to bird species (*23*), and the solidline represents the result of a high-profile drag coefficient (CD_pro_ = 0.075) based on the reconstruction of pterosaur wings (*22*, *57*). In (A), the maximum glide ratios of each species are shown on the right side of species names. Points represent the horizontal speed and sinking speed at the maximum glide ratio of each species. (B) shows a circling envelope. The left end of the curve is for a bank angle of 45 degrees. The smaller the bank angle, the larger the circle radius. The linear wingspan reduction is assumed. The lift coefficient of circling envelope 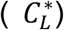 is the lift coefficient at the minimum sinking speed.

With respect to the soaring up phase, all of the extinct giant flyers, except for *Quetzalcoatlus*, had performances equivalent to or better than the extant species (Fig. 5B). As shown previously (*4*), the circling up performance of *Argentavis magnificens* was comparable to that of the California condor, one of the largest living thermal soarers. The performance of *Pteranodon* was comparable to that of living thermal soarers, irrespective of the value of the assumed profile drag coefficient. At low drag coefficients, the minimum sinking speed of *Pteranodon* was similar to that of the white stork, and at high drag coefficients, the performance was similar to that of the California condor and the *Argentavis magnificens*, although *Pteranodon* required a slightly larger turning radius than these species. The thermal soaring ability of *Pelagornis sandersi* when a light mass was assumed (21.9 kg) was outstanding. It outperformed several extant thermal soaring species in soaring up ability, and was even comparable to the magnificent frigatebirds, the champion of thermal soaring among extant species. Even with a heavier body mass estimate (40.1 kg), *Pelagornis sandersi* still outperformed or was comparable to several other species.

Among the four extinct giant animals investigated in this study, the soaring up performance of *Quetzalcoatlus* was exceptionally low. It required the strongest upward wind speed and the widest circle radius. Its performance was even lower than that of the kori bustard, one of the heaviest volant extant bird species, which spends most of its time on land and only fly in emergencies, such as when under predation risk.

It also demonstrates that motor gliders perform very differently from birds and pterosaurs. While motor gliders can achieve a small sinking speed, they require a larger circle radius than animals, reflecting the fact that they have a higher wing loading than *Quetzalcoatlus* and lower maximum lift coefficient (1.3, assumed in a previous study (*33*)). In contrast, due to their low drag, gliders outperform all birds and pterosaurs in gliding performance. For example, compared with *Quetzalcoatlus*, which has a similar wing span and weight, the glider has a maximum glide rate two to three times higher. This means that while a *Quetzalcoatls* can travel 8 or 14 m horizontally during 1 m descent, a glider can travel 28 m (Fig. 5A).

## Discussion

Although several previous studies have investigated the soaring performance of extinct species, there have been several evaluation gaps. In the present study, we filled these gaps using physical models of soaring birds. We computed and compared the dynamic and thermal soaring performances and the required wind conditions for soaring of four extinct giant flyers with those of extant dynamic and thermal soaring species, which enabled us to examine the soaring style of extinct giants from multiple perspectives.

Our results indicate that *Argentavis magnificens* and *Pteranodon* were thermal soarers, confirming previous studies (*4*, *22*). However, our results also indicate that *Quetzalcoatlus* could not efficiently perform dynamic or thermal soaring. In addition, although *Pelagornis sandersi* was considered a dynamic soaring species in a previous study (*1*), our results suggest that it was a thermal soaring bird. We discuss our results in detail for *Quetzalcoatlus* and *Pelagornis sandersi* and then describe future issues that need to be addressed for a better understanding of the soaring styles of extinct giant species.

### Quetzalcoatlus

There has been a heated debate about the flight capability of *Quetzalcoatlus*. The focal issue has been whether or not *Quetzalcoatlus* could take off. Researchers are divided between the opinion that it was too heavy to take off (*17*, *21*, *34*) and the opinion that it was able to take off by using quadrupedal launching, like some bats (*13*, *18*). In addition, detailed observations of fossils are also presented as evidence that the giant azhdarchids, including *Quetzalcoatlus*, were capable of flight; for example, a huge deltopectoral crest on their humeri, which would have anchored muscles for flapping flight (*35*).

Although there is some debate as to whether or not giant pterosaurs could have taken off, it has been widely accepted that if they were able to take off their primary mode of travel would have been thermal soaring rather than flapping flight (*13*). Witton and Habib applied a model of bird flap-gliding flight (*23*) to *Quetzalcoatlus* and found that the flapping flight of this species required anaerobic movement and was difficult to sustain for a long period; therefore, it must have relied on wind energy for long-distance travel (*13*). Based on a comparison of morphology between birds, bats, and pterosaurs with principal component analysis (PCA), these authors also concluded that *Quetzalcoatlus* used thermal soaring (and that *Pteranodon* used dynamic soaring) (*5*, *13*).

Our results revealed that *Quetzalcoatlus* had a poor ability to use thermals to ascend. It required a larger circle radius and stronger updraft than the terrestrial kori bustard, let alone the species that use thermal soaring. Whether *Quetzalcoatlus*, with the soaring performance shown in Fig. 5, could routinely travel long distances by thermal soaring is beyond the scope of this study because of the need to examine in detail the wind conditions in their habitat at the time they lived. However, the results of this study alone suggest that *Quetzalcoatlus* performed poorly at thermal soaring compared with modern and other extinct species, and that the wind conditions under which thermal soaring was possible were limited. This poor thermal soaring performance was due to the large wing loading associated with the large body size. As shown in the **Materials and Methods**, the circle radius was proportional to the wing loading to the power of one half (eq. 19), and the descent speed during the turn was also proportional to the wing loading to the power of one half if the effect of the organism’s wing length adjustment was ignored (eqs. 18 and 20, and see also (*32*)). Since the wing loading was approximately proportional to body size, a giant *Quetzalcoatlus* required thermals with a wider radius and stronger updraft for thermal soaring. The wing loading also explains why the results of the present study are not consistent with the claims of previous studies that *Quetzalcoatlus* was adapted to thermal soaring (*5*, *13*). In the previous study, soaring ability was assessed from two variables related to wing loading and aspect ratio with PCA (second and third principal components obtained by PCA on the logarithms of body weight, wing area, and wingspan. The first principal component is roughly related to body size. See (*36*, *37*) for details of the method and data). These variables are not exactly equal to wing loading and aspect ratio. In particular, the size-dependence has been removed from the principal component related to wing loading. However, thermal soaring performance is inevitably size-dependent and, therefore, caution should be taken when evaluating soaring ability with PCA. When evaluating soaring performance from animal morphology, using performance and wind requirements calculated from morphology based on the laws of physics (as conducted in this study and (*15*, *32*)) are more accurate, as taking an inappropriate combination of morphology as a variable may lead to erroneous results.

Anatomical studies of the azhdarchid pterosaurs have reported that their skeletal structure shows adaptations to terrestrial walking and suggested that they were terrestrial foragers (*38*, *39*). Furthermore, a recent phylogenetic analysis showed that the azhdarchoid pterosaurs differed from other pterosaurs in that they had evolved in a manner that increased the cost of transport for flapping flight and the sinking speed of gliding (*15*). Taking into account the adaptations for walking (*38*, *39*), the humeri feature indicating flapping flight capability (*35*) but not sustainable flapping flight (*13*), the phylogenetic tendency of decreasing flight efficiency (*15*), and the low thermal soaring ability shown here, we suggest that the flight styles of *Quetzalcoatlus* and other similar-sized azhdarchid *s*pecies were similar to those of the bustard or ground hornbill that spend most of their time on land and rarely fly. Alternatively, they may have routinely flown in early stages of their life history, but as they matured, wing loading would increase, and they would spend most of their time on land.

### Pelagornis sandersi

As *Pelagornis sandersi* was found close to the coast, this species is thought to have lived an oceanic existence by soaring over the sea (*1*). Previously, it has been reported that *Pelagornis sandersi* was a dynamic soarer like the albatross, rather than a thermal soarer like frigatebirds (*1*). However, we argue that this species is highly adapted for thermal soaring rather than dynamic soaring. The conclusion of the previous study was based on the glide polars of *Pelagornis sandersi*, which were more similar to those of the wandering albatrosses than those of frigatebirds; glide performance was the only criterion used to evaluate its soaring style. In this study, we quantified other performances and the required wind conditions, which enabled us to evaluate the soaring style of *Pelagornis sandersi* from multiple perspectives.

Our results indicated that the dynamic soaring performance of *Pelagornis sandersi* was generally inferior to that of extant dynamic soaring species, although there were substantial variations depending on wind conditions and morphology estimates. One of the factors contributing to the poor dynamic soaring ability of this species was an inability to efficiently exploit the wind speed gradient due to long wings limiting the height above sea level at which the bird could fly. This effect could not be assessed using the glide polars alone.

Conversely, the thermal soaring ability of *Pelagornis sandersi* was outstanding regardless of the morphology estimates used. The thermal soaring ability of an animal is largely dependent on its wing loading, as previously discussed. Therefore, the reason why *Pelagornis sandersi* showed such a high performance is because of its low wing loading despite its huge size. Considering that *Pelagornis sandersi* was found close to the coast, this species is expected to have been well adapted to capture weak updrafts above the sea by using thermal soaring, and that it was able to stay aloft for a long period of time with limited flapping and traveled long distances, similar to frigatebirds (*40*).

### Future issues

In this section, we discuss some of the simplifications used in this study and issues that we believe need to be addressed in the future.

The first issue is flight stability. In this study, a steady wind environment was assumed, but actual wind environments fluctuate. In such a fluctuating real-world environment, stability is an important factor that determines the success or failure of flight (*41*, *42*). To simplify our calculations, we did not address stability, but it is important to examine the flight stability of these extinct and extant birds using more detailed morphological information in the future.

The next issue is that the actual wind environment experienced by animals is still largely unknown. For dynamic soaring, the specific form of the wind speed gradient experienced by birds is unknown—for example, whether there is a logarithmic or sigmoidal gust in the shadows of waves—and information on the height and thickness of the wind speed gradient is not yet known (See **Wind gradient models** in **Materials and Methods**) (*31*, *43*) For this reason, we evaluated performance under various wind conditions (Fig. 3). For thermal soaring, it is also unknown how much updraft animals experience at a given circle radius or the distance between thermals. Recent advances in tracking technology have made it possible to record details of the motion of birds in dynamic and thermal soaring (*44*–*48*). We believe that these data will provide information that will aid in understanding our results and improving our model, such as the real wind environment experienced by animals (*48*–*52*). In addition, it is also important to consider the paleoenvironmental aspects of the wind environment at the time of the extinct species’ inhabitation. For example, we showed that *Quetzalcoatlus* had a lower thermal soaring capacity than the extinct species. Paleoclimatic estimates may help us to understand whether the species had a favorable wind environment that allowed it to use thermal soaring as its primary mode of transport, even with their poor soaring up ability.

Third, it should be noted that in our model of dynamic soaring, the maximum speed the animal could achieve was very high. Depending on the shape of the wind speed gradient assumed in our model, the animals reached maximum speeds of over 100 km/h even at realistic wind speeds of 5–10 m/s (Fig. 4), however, the average speeds reported using GPS for albatrosses and shearwaters were about 30–55 km/h (*51*, *53*). There are two potential reasons for the discrepancy between this model and reality. The first reason is that the actual shape of the wind speed gradient is likely to be more gradual than that assumed in the present study. For example, it can be seen from our results in a sigmoidal wind gradient (Fig. 4 B and C) that the gentler the wind speed variation (i.e., the larger the *δ*), the slower the maximum travel speed. Although little is known about the shape of the wind gradients experienced by actual birds as mentioned in the previous paragraph, the actual wind speed gradient may be closer to a sigmoidal form, for example, with a larger value of *δ* than assumed. The second potential reason is that the cost of rolling was not taken into account. Dynamic soaring puts the wings at risk of breaking under the influence of the high moment of force. Accordingly, the wing strength may limit the speed of change in bank angle and the resulting flight speed of animals. This would be particularly detrimental for species with long wingspans, such as the wandering albatross and the extinct giants. Indeed, reinforcing wing strength is also an important issue when designing an unmanned aerial vehicles (UAV) for dynamic soaring (*54*). In view of the above two points, the dynamic soaring performance and the required wind conditions obtained from the present numerical calculation should not be taken literally by the values themselves. However, since all species were evaluated under the same assumptions, the relative values are meaningful indicators for the purpose of estimating soaring style by inter-species comparison, as was done in this study. A more refined analysis incorporating constraints on body rolling speed will be an interesting challenge in the future. For this purpose, actual measurements of bank angles in dynamic soaring birds and assessment of wing bone strength in extant and extinct species will provide important information.

In summary, we clarified what needs to be examined and what further information is required, which is an important outcome of this research. Understanding the soaring performance of extinct animals is an interdisciplinary issue. It is therefore essential to have a place for objective discussion between researchers with different backgrounds to bring their knowledge together, but such a platform seems to have been lacking in the past. In our approach, i.e., the physical model and the framework for comprehensively evaluating soaring performance, we have tried to clarify what assumptions were needed regarding the mechanics and the morphology of the animals. On the basis of our theoretical framework, it should therefore be easy for experts from various disciplines, including paleontologists, engineers, and ornithologists, to examine the validity of the assumptions, examine new information, such as updated morphological data, and improve the model. We hope that our theoretical framework presented in this study will inspire researchers from various disciplines to work together to understand the soaring performance of extinct animals.

## Materials and Methods

### Quantification of soaring styles in previous studies

This section reviews previous studies on the soaring performance of extinct giant animals. In particular, we focus on which indices were quantitatively evaluated for each species.

#### Pelagornis sandersi

*Pelagornis sandersi* is predicted to be a dynamic soarer rather than a thermal soarer as its glide polar (and glide ratio that can be derived from its glide polar) is more similar to those of living dynamic soarers than those of living thermal soarers (*1*). However, this means the understanding of this species’ soaring style has been based on just one metric, that is, its glide ratio (Table 2). Hence, evaluating other metrics of this species could provide a more accurate estimate of the soaring style of this species. A previous study cautiously calculated the glide polars of *Pelagornis sandersi* for 24 combinations of estimates (body mass = 21.8 and 40.1 kg, wingspan = 6.06, 6.13, 6.40, and 7.38, and aspect ratio = 13,14, and 15) to deal with morphological uncertainty. Hence, we also employed these estimates in this study.

#### Argentavis magnificens

*Argentavis magnificens* is expected to be a thermal soarer. A previous study reported that the thermal soaring performance and required wind conditions of this species were comparable to living thermal soaring species based on glide polars and circling envelopes (*4*). This result is consistent with the fact that an *Argentavis* specimen was found on the foothills and pampas of Argentina, far from coastlines (*4*).

#### *Pteranodon* and *Quetzalcoatlus*

Although assessments of the soaring abilities of *Pteranodon* and *Quetzalcoatlus* have been a long-standing issue, there is still lack of a comprehensive understanding of their soaring style due to several uncertainties in the estimates of their morphology, especially because of the significant changes in weight estimates around 2010. Previously, it was estimated that *Pteranodon* had a wingspan of around 7 m and a body mass of 16 kg, while *Quetzalcoatlus* had a wingspan of around 11 m and a body mass of 50–70 kg. Based on these estimates, previous studies argued that they were adapted to thermal soaring (*55*, *56*) and others argued that they could also employ dynamic soaring (*21*). Around 2010, however, several studies with different approaches suggested that pterosaurs were much heavier than previously expected (*5*, *17*, *29*). For example, Witton estimated that *Pteranodon* was 36 kg with a 5 m wingspan, and *Quetzalcoatlus* was 259 kg with a 10 m wingspan (*5*). Other studies also reported similar estimated values (*17*, *29*). Few studies have quantified the soaring performance of these species based on these new heavy body mass estimates. For example, Witton and Habib argued that *Pteranodon* was a dynamic soarer and *Quetzalcoatlus* was a thermal soarer by comparing the second and third principal components calculated by PCA of the logarithms of weight, wing area, and wingspan of these species with those of extant flying bird species. (*5*, *13*, *37*). Conversely, a recent study quantified the cost of transport and sinking speeds during gliding in pterosaur species and showed that azhdarchoid pterosaurs, including *Quetzalcoatlus*, had lower flight efficiency than the other pterosaurs (*15*). Despite these studies, the performance and wind requirements of dynamic and thermal soaring in these species have not been comprehensively quantified. Furthermore, pterosaurs have a wing morphology that is completely different from that of birds and bats. Some studies reported that the wings of pterosaurs would have been associated with high-profile drag (drag stemming from the wings) (*22*, *57*). Palmer experimentally measured profile drag in a wind tunnel experiment using reconstructed pterosaur wings (*22*). With the experimentally derived profile drag, glide polars of *Pteranodon* with various body mass estimates including the recent heavy estimate were determined. Palmer concluded that *Pteranodon* adopted thermal soaring with a slow flight speed. Evidence of the pterosaur’s slow flight was further reinforced by Palmer’s subsequent work that quantified the strength of the pterosaur’s wing membranes using a physical approach, with the result that their wing membranes were not suited to fast flight (*58*).

### Models

The dynamics of soaring animals are described using the equations of motion (EOM). We first describe the EOM and parameters therein. Using the EOM, we can calculate soaring performances and the wind speeds required for sustainable soaring. We then describe the calculation procedure for dynamic soaring and thermal soaring, respectively.

#### Aerodynamic forces and parameters

We regard an animal as a point of mass. The dynamics of the animal’s three-dimensional position ***X***(*t*) and velocity ***V***(*t*) are represented by the following EOMs:

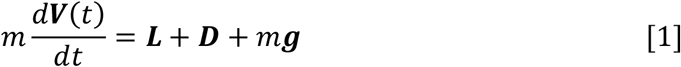

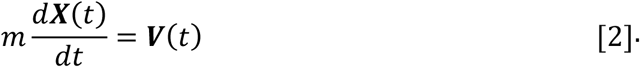

When an animal is soaring, three forces—gravitation (*m***g**), lift force (***L***), and drag force (***D***)—act on it. Gravitation *m**g*** is a product of the constant of gravitation (***g***) and mass of the bird (*m* kg), and its direction is towards the ground. The direction of the lift force ***L*** is dorsal and perpendicular to the air velocity. Drag force ***D*** is against the air velocity. For the analysis of dynamic soaring, we assumed that wind blow is along the y-axis and represent these EOMs in a different way by transforming the ground velocity to pitch *γ*, yaw *ψ*, bank angle *ϕ*, and airspeed *V* using the following equations (*31*):

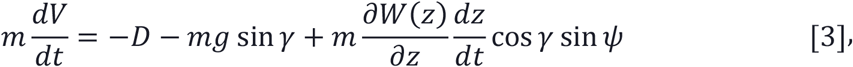

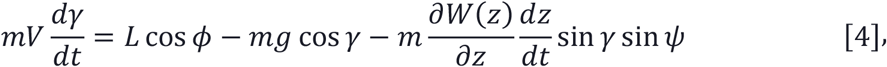

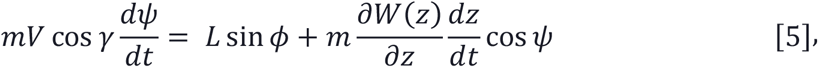

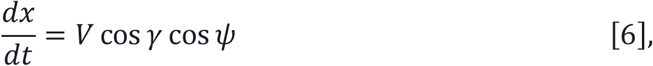

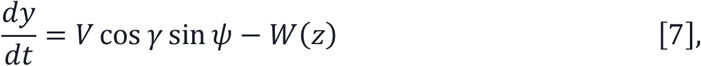

and

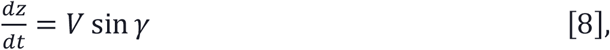

where *W*(*z*) represents the wind gradient. A specific form is provided in the latter subsection. *L* represents the strength of the lift force, and *D* represents that of the drag force. The aerodynamic theory asserts that these values are

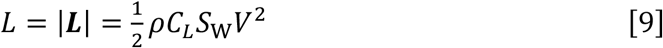

and

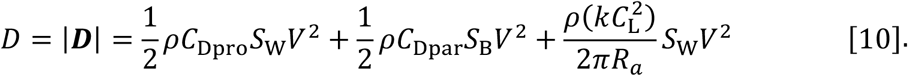

Here, *ρ* is air density and was set to *ρ* = 1.23 kg m^−3^ (*15*). This is the International Standard Atmosphere value for sea level at 15 °C expressed as 3 significant digits (*1*). *C*_L_ represents the lift coefficient. *S*w represents the wing area. The drag is composed of three terms. The first term is the profile drag that stems from friction on the wing. *C*_Dpro_ is the profile drag coefficient. For birds, a *C*_Dpro_ of 0.014 was employed following previous studies (*1*, *23*). However, based on the reconstruction of pterosaur wings and a wind tunnel experiment, some studies have argued that the pterosaur profile drag coefficient is much higher (approximately 0.05–0.1) (*22*, *57*). Hence, for the profile drag of *Pteranodon* and *Quetzalcoatlus*, we explored two cases: a bird-like low-profile drag of 0.014 and an experimentally based high-profile drag of 0.075 (we note that, although the experiments in the previous study assumed a pterosaur with a wing span of approximately 6 m (*22*), we assumed that the result would not vary significantly with the size of *Quetzalcoatlus*). The second term is the parasite drag stemming from friction on the body, where the *C_Dpar_* is the parasite drag coefficient, and *S*_B_ is the body frontal area. We used the following recently recommended formula

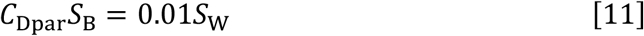

on the practical basis that neither *C*_Dpar_ nor *S*_B_ is exactly known (*32*). The third term is the induced drag that stems from the lift force. *R*_a_ represents the aspect ratio (*R*_a_ = *b*^2^ / *S*w, where b is the wingspan). *k* is the induced drag factor; we set *k* to 1.1, as in previous studies (*1*, *13*, *23*). The lift coefficient has a maximum value; for birds, we set *C*_L_ to ≤ 1.8 (*23*). As the aerodynamic properties of pterosaurs can differ from those of birds, and the wind tunnel experiment indicated that *C*_L_ could reach more than 2.0 (*22*), we set the pterosaurs’ lift coefficient to ≤ 2.2.

The remaining parameters in the EOMs are body mass (*m*), wingspan (*b*), and wing area (*S*w). For these morphological parameters of extant birds, we used values reported in previous studies. For *Pelagornis sandersi*, we used 24 combinations of estimates proposed in a previous study (*1*). For *Argentavis magnificens*, we used the estimates in (*4*). For *Pteranodon* and *Quetzalcoatlus*, we used the recent estimates for heavy pterosaurs, as reported in (*5*). These values are shown in Table 1.

The EOMs include variables that soaring animals can control, that is, bank angle *ϕ* (t) and lift coefficient *C*_L_. Although these variables are time-dependent, for simplicity, we assumed that the animals keep their lift coefficients at a constant value. Hence, using a time series for bank angle, a constant value of *C*_L_, and values of parameters, the dynamics of the soaring animals were determined with EOM.

#### Quantification of the dynamic soaring performance and the required minimum wind speed

### Wind gradient model

We explored two types of wind gradients. The first was the logarithmic model represented as

[Logarithmic wind gradient model]

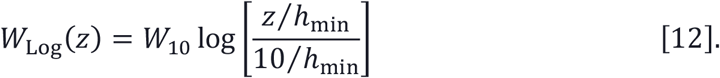

This function is defined at *z* > *h*_min_. We set *h*_min_ to 0.03 [m], following a previous study (*28*). *W*_10_ is the wind speed at height *z* = 10 m. This model is deemed to be a good model of the average wind field in the first 20 m above the sea surface, assuming a flat sea surface, and has been a popular approach in dynamic soaring modeling. However, recent studies argued that the real sea surface is not flat, and wind separations in ocean waves may occur more often than expected (*43*). To describe wind-separation-like wind profiles, a sigmoidal model has been proposed (*31*, *59*). We also employed the sigmoidal wind model with a minor change, represented as

[Sigmoidal wind gradient model]

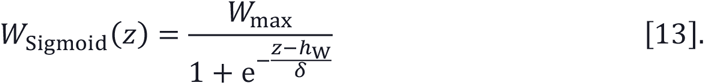

*h*_w_ determines the height of wind separation, as shown in Fig. 3. In this study, we set *h*_w_ to 1, 3, and 5. δ is the thickness parameter. The wind speed changes with height (|*z* − *h_w_*| ≲ 3*δ* m). In a previous study, the wind shear thickness was speculated as approximately 1.5–7 m. Here, we set *δ* to 3/6 with a steep wind change, and 7/6 with a gentler change (Fig. 3).

### Formulation to numerical optimization

The numerical computation of dynamic soaring performance and minimum wind speed boiled down to the restricted optimization problem (*60*). That is, a mathematical problem to find the values of (i) a certain variable ***Y*** that maximizes (ii) an objective function *f*(***Y***), satisfying (iii) equalities ***h***(***Y***) = 0 and (iv) inequalities ***g***(***Y***) ≤ 0. In the following, we describe the variables, object functions, equalities, and inequalities for dynamic soaring.

#### (i) Variables

The dynamics of dynamic soaring animals are described by the 3D position (*x*(t), *y*(t), *z*(t)), pitch angle *γ*(*t*), yaw angle *ψ*(*t*), airspeed *V*(*t*), bank angle *ϕ*(*t*), lift coefficient *C*L, and the period of one dynamic soaring cycle *τ*. Among these variables, 3D position, pitch, yaw, bank, and airspeed are functions of time *t* (0 ≤ *t* ≤ *τ*). Optimization problems that include functions as vari**4** ables are difficult to be directly solved. Therefore, we employed a collocation approach (*30*, *31*). The collocation approach discretizes the variables in time, such as *X*(t) (0 ≤ *t* ≤ *τ*) to variables *X_i_* = *X*((*i*−1)*N*/*τ*) (*i*= 1, *N*), and converts the EOM to the equalities between those discretized variables. Hereafter, we use *X*_1:*N*_ = {*X*_1_, *X*_2_,…, *X_N_*}. In this study, we set the number of discretization points to *N* = 51 in order to perform computations with reasonable accuracy within a reasonable amount of time. Accordingly, the variables of this optimization problem are position *x*_1:*N*_, *y*_1:*N*_, *z*_1:*N*_, pitch angle *γ*_1:*N*_, yaw angle *ψ*_1:*N*_, airspeed *V*_1:*N*_, bank angle *ϕ*_1:*N*_, lift coefficient *C*L, and a period of one soaring cycle *τ*. In addition, when computing the minimum wind speed required for sustainable dynamic soaring, *W*_10_ (log model) or *W*_max_ (sigmoid model) were also treated as variables. Hence, the total number of variables were 7 × 51 + 2 (+1 [when computing the minimum wind speed])= 359 (or 360).

#### (ii) Object function

First, we computed (1) the minimum wind speed required for sustainable dynamic soaring for each wind gradient model. As the objective function to minimize, we set *W*_10_ for the logarithmic model and *W*_max_ for the sigmoidal model. Then, we computed (2) the maximum speed averaged over one dynamic soaring cycle by maximizing the object function to 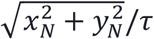. Finally, we computed (3) the maximum upwind speed averaged over one dynamic soaring cycle by maximizing the object function *y_N_*/*τ*. With respect to the maximum speed and maximum windward speed, we computed these values for different wind speeds, that is, from the minimum required wind speed of the species to the highest minimum required wind speed among the examined species (i.e., *Quetzalcoatlus*) + 2 m/s. In this wind speed range, the maximum speed reached an unrealistically high value and/or the optimization calculation did not converge for some species. Thus, we stopped the computation of the maximum speed at the wind speed where the maximum speed exceeded 40 m/s (144 km/h).

#### (iii) Equalities

The first equalities to be fulfilled for dynamic soaring animals are given in Eqs. 3–8. The collocation approach converts the EOM into the equalities between the variables listed in the above section. As the number of original EOM was six and the number of discretization was 51, the EOM were converted into 6 × 51 = 306 equalities (see (*30*, *31*) for the specific representations of these equalities).

The second type of equalities to be fulfilled were periodic boundary conditions of dynamic soaring: at the beginning and end of one dynamic soaring cycle, the state of the animal (i.e., pitch, yaw, airspeed, bank, height) is the same, represented as

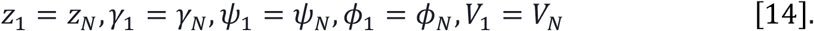

#### (iv) Inequalities

First, we assumed that there was a maximum limit of physical load on the animal. This is because dynamic soaring entails dynamic maneuvering, which results in a corresponding acceleration. We employed the approach of a previous study (*28*) that restricted the load factor (*L*/*mg*) to less than 3,

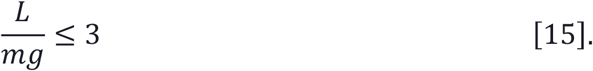

The second inequality was an important modification of the previous models. The height of the animal’s wingtip above the sea surface (*z*_wing_) was calculated and represented as

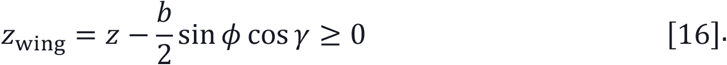

Previous studies discarded the existence of the sea surface (*31*) or restricted birds to only flying higher than a given height (1.5 m) from the sea surface (*28*). However, the height a bird can fly depends on the wing length and the bank angle (e.g., with a shorter wing length and a lower bank angle, a bird can fly at a lower height). When dynamic soaring birds fly, they adjust their wingtips close to, but avoid touching, the sea surface (Fig. 3D). Dynamic soaring birds can exploit more flight energy when they pass through stronger wind speed differences. As the wind speed difference is strong close to the sea surface, how close to the sea surface a bird can fly is crucial for dynamic soaring birds. Accordingly, long wings may restrict the minimum height at which the bird can fly and disturb efficient dynamic soaring. Hence, considering the effect of wings is crucial for evaluating dynamic soaring performances.

Third, we assumed that the height of the animal was higher than 0.5 m, that is,

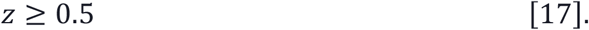

The optimization problem described here is a restricted non-linear optimization problem. We used the SQP method to solve the problem with the ‘fmincon’ function in MATLAB® Ver R2019a.

#### Quantification of the thermal soaring performance and the required upward wind speed

For the computation of glide polars and circling envelopes, we followed the same procedure as the *Flight* software developed for evaluating bird flight performance, as described in (*23*). In the following, we outline the procedure and parameters employed in this study.

First, before computing the glide polars, we determined how gliding animals adjust their wingspan with respect to their airspeed. Three wingspan reduction ways (linear wingspan reduction, wing-drag minimizing wingspan reduction, and fixed wingspan) are presented in (*23*). We conducted calculations using each reduction ways and found no substantial differences in the results (see Fig. 5 and SI Appendix). We show the results using the *Flight’s* default settings, which have been used in previous studies (*1*, *13*). These assume that wingspan, wing area, and thus aspect ratio linearly decrease with factor *β* = (*B*_stop_ - *V*/*VS*)/(*B*_stop_ − 1); i.e., we replaced *b*, *S*w, and *R*_a_ with *βb*, *βS*w, and *βR*_a_, respectively. In this equation, *V_S_* is the stall speed, the airspeed of the animal at the highest lift coefficient (i.e., 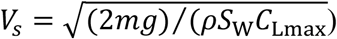); and *B*_stop_ is a constant that determines the degree of wing reduction. For birds, we set *B*_stop_ to 5, the default value in *Flight*. For pterosaurs, we set *B*_stop_ to 6, following a previous study (*13*). Then, the glide polars were derived from the EOM, setting bank angle to 0, assuming that the pitch angle was small enough (*γ* ≪ 1) and considering the gliding animal was at kinematical equilibrium (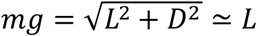, sin *γ* = *D*/*L* ≃ *L*/*mg*). Sinking speed was represented as a function of airspeed *V* (*23*),

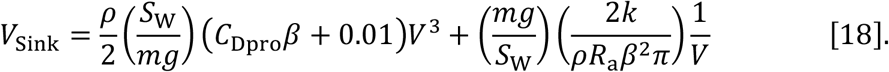

This relation gives a glide polar. Note that we used eq. 11 (*C*_Dpar_ *S*_B_ = 0.01*S*w; in this equation, *S*w was not replaced with *βS*w) to derive the above equation. The horizontal speed is 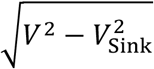. Thus, the maximum glide ratio is the maximum value of 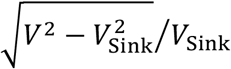.

The circling envelope is given with the radius of the circle (*r*), and the sinking speed in a circling glide (*V*_Sink,Circle_) represented by the bank angle (*ϕ*),

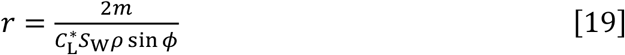

and

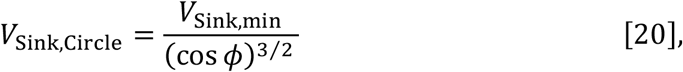

where *V*_Sink,min_ is the minimum sinking speed in straight flight calculated from eq. 18. The 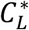 is the lift coefficient and two different values are often employed. The first one is that 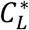 is the maximum lift coefficient (*33*, *61*). The other one is that 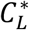 is the lift coefficient at the minimum sinking speed (*23*, *62*). The former gives the lowest sinking speed in circling a given radius, whereas the animal is flying at a stall speed. The latter gives a greater sinking speed in circling a given radius than the former, but the airspeed is greater than the stall speed (unless the airspeed at the minimum sinking speed is the same as the stall speed). As the bank approaches vertical (*ϕ* → 90°), the radius approaches a non-zero value called the limiting radius of turn (*23*). The former definition gives the minimum value of limiting radius. As there was no substantial difference in either case, the results for the latter definition are presented in Fig, 5B (see Appendix SI for the former results). The limiting radius, minimum sinking speed in straight gliding, maximum glide ratio, and horizontal speed at maximum glide ratio are shown in Appendix SI for the three wing reduction methods and 2 circling envelope definition (i.e., 6 cases). For ASK14, the measured glide polar (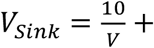 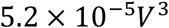) and the circling envelope estimated from them, reported in a previous study (*33*), are presented.

## Supplementary Materials

Fig. S1. Glide polars and circling envelopes where 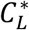 is the lift coefficient at the minimum sinking speed assuming a wing-drag minimizing wingspan reduction.

Fig. S2. Glide polars and circling envelopes where 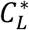 is the lift coefficient at the minimum sinking speed assuming a fixed wingspan.

Fig. S3. Glide polars and circling envelopes where 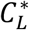 is the maximum lift coefficient assuming a linear wingspan reduction.

Fig. S4. Glide polars and circling envelopes where 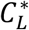 is at the maximum lift coefficient.

Fig. S5. Glide polars and circling envelopes where 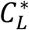 is at the maximum lift coefficient.

Table S1. Thermal soaring performances where 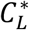 is at the minimum sinking speed assuming linear wingspan reduction

Table S2. Thermal soaring performances where 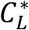 is at the minimum sinking speed assuming a wing-drag minimizing wingspan.

Table S3. Thermal soaring performances where 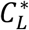 is at the minimum sinking speed assuming a fixed wingspan.

Table S4. Thermal soaring performances where 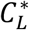 is the maximum lift coefficient assuming linear wingspan reduction.

Table S5. Thermal soaring performances where 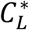 is the maximum lift coefficient assuming a wing-drag minimizing wingspan.

Table S6. Thermal soaring performances where 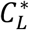 is the maximum lift coefficient assuming a fixed wingspan.

## Acknowledgments

The authors thank Chihiro Kinoshita for illustrating Fig. 1 and Shin-ichi Fujiwara, Sophia Friesen, and Michael Le Page for reading the draft or preprint of this paper and for providing valuable comments. This study was financially supported by Grants-in-Aid for Scientific Research from the Japan Society for the Promotion of Science (16H06541, 16H01769), and JST CREST Grant Number JPMJCR1685, Japan. The authors also received financial support from JSPS as part of an Overseas Research Fellowship program.

## Author contributions

YG, KY, HW, and KS designed the study; YG conducted the computation; all authors participated in the writing of the manuscript (YG wrote the first draft).

## References

1. D. T. Ksepka, Flight performance of the largest volant bird. Proc. Natl. Acad. Sci. 111, 10624–10629 (2014).

2. K. E. Campbell Jr, E. P. Tonni, Preliminary observations on the paleobiology and evolution of teratorns (Aves: Teratornithidae). J. Vertebr. Paleontol. 1, 265–272 (1981).

3. K. E. Campbell Jr, E. P. Tonni, Size and locomotion in teratorns (Aves: Teratornithidae). Auk. 100, 390–403 (1983).

4. S. Chatterjee, R. J. Templin, K. E. Campbell, The aerodynamics of Argentavis, the world’s largest flying bird from the Miocene of Argentina. Proc. Natl. Acad. Sci. 104, 12398–12403 (2007).

5. M. P. Witton, A new approach to determining pterosaur body mass and its implications for pterosaur flight. Zitteliana, 143–158 (2008).

6. E. Frey, D. M. Martill, A reappraisal of Arambourgiania (Pterosauria, Pterodactyloidea): one of the world’s largest flying animals. Neues Jahrb. für Geol. und Paläontologie-Abhandlungen, 221–247 (1996).

7. E. Buffetaut, D. Grigorescu, Z. Csiki, Giant azhdarchid pterosaurs from the terminal Cretaceous of Transylvania (western Romania). Geol. Soc. London, Spec. Publ. 217, 91–104 (2003).

8. D. W. E. Hone, M. B. Habib, F. Therrien, Cryodrakon boreas, gen. et sp. nov., a Late Cretaceous Canadian Azhdarchid Pterosaur. J. Vertebr. Paleontol. 39, e1649681 (2019).

9. M. Vremir, G. Dyke, Z. Csiki‐Sava, D. Grigorescu, E. Buffetaut, Partial mandible of a giant pterosaur from the uppermost Cretaceous (Maastrichtian) of the Haţeg Basin, Romania. Lethaia. 51, 493–503 (2018).

10. E. H. Hankin, D. M. S. Watson, On the flight of pterodactyls. Aeronaut. J. 18, 324–335 (1914).

11. G. Mayr, V. L. De Pietri, L. Love, A. Mannering, R. P. Scofield, Oldest, smallest and phylogenetically most basal pelagornithid, from the early Paleocene of New Zealand, sheds light on the evolutionary history of the largest flying birds. Pap. Palaeontol. (2019).

12. P. A. Kloess, A. W. Poust, T. A. Stidham, Earliest fossils of giant-sized bony-toothed birds (Aves: Pelagornithidae) from the Eocene of Seymour Island, Antarctica. Sci. Rep. 10, 1–11 (2020).

13. M. P. Witton, M. B. Habib, On the size and flight diversity of giant pterosaurs, the use of birds as pterosaur analogues and comments on pterosaur flightlessness. PLoS One. 5, e13982 (2010).

14. R. B. J. Benson, R. A. Frigot, A. Goswami, B. Andres, R. J. Butler, Competition and constraint drove Cope’s rule in the evolution of giant flying reptiles. Nat. Commun. 5, 1–8 (2014).

15. C. Venditti, J. Baker, M. J. Benton, A. Meade, S. Humphries, 150 million years of sustained increase in pterosaur flight efficiency. Nature (2020), doi:10.1038/s41586-020-2858-8.

16. T. J. Pedley, in International Symposium on Scale Effects in Animal Locomotion (1975: Cambridge University) (Academic Press, 1977).

17. K. Sato, K. Q. Sakamoto, Y. Watanuki, A. Takahashi, N. Katsumata, C.-A. Bost, H. Weimerskirch, Scaling of soaring seabirds and implications for flight abilities of giant pterosaurs. PLoS One. 4, e5400 (2009).

18. M. B. Habib, Comparative evidence for quadrupedal launch in pterosaurs. Zitteliana, 159–166 (2008).

19. A. E. R. Cannell, Too big to fly? An engineering evaluation of the fossil biology of the giant birds of the Miocene in relation to their flight limitations, constraining the minimum air pressure at about 1.3 bar. Anim. Biol. 1, 1–20 (2020).

20. H. J. Williams, E. L. C. Shepard, M. D. Holton, P. A. E. Alarcón, R. P. Wilson, S. A. Lambertucci, Physical limits of flight performance in the heaviest soaring bird. Proc. Natl. Acad. Sci. 117, 17884–17890 (2020).

21. S. Chatterjee, R. J. Templin, Posture, locomotion, and paleoecology of pterosaurs (Geological Society of America, 2004), vol. 376.

22. C. Palmer, Flight in slow motion: aerodynamics of the pterosaur wing. Proc. R. Soc. B Biol. Sci. 278, 1881–1885 (2011).

23. C. J. Pennycuick, Modelling the flying bird (Elsevier, 2008), vol. 5.

24. C. J. Pennycuick, Thermal soaring compared in three dissimilar tropical bird species, Fregata magnificens, Pelecanus occidentals and Coragyps atratus. J. Exp. Biol. 102, 307–325 (1983).

25. G. P. Sachs, in AIAA Scitech 2019 Forum (2019), p. 568.

26. P. L. Richardson, Upwind dynamic soaring of albatrosses and UAVs. Prog. Oceanogr. 130, 146–156 (2015).

27. G. Sachs, In-flight measurement of upwind dynamic soaring in albatrosses. Prog. Oceanogr. 142, 47–57 (2016).

28. G. Sachs, Minimum shear wind strength required for dynamic soaring of albatrosses. Ibis (Lond. 1859). 147, 1–10 (2005).

29. D. M. Henderson, Pterosaur body mass estimates from three-dimensional mathematical slicing. J. Vertebr. Paleontol. 30, 768–785 (2010).

30. Y. J. Zhao, Optimal patterns of glider dynamic soaring. Optim. Control Appl. methods. 25, 67–89 (2004).

31. G. D. Bousquet, M. S. Triantafyllou, J.-J. E. Slotine, Optimal dynamic soaring consists of successive shallow arcs. J. R. Soc. Interface. 14, 20170496 (2017).

32. G. K. Taylor, A. L. R. Thomas, Evolutionary biomechanics: selection, phylogeny, and constraint (Oxford Series in Ecology and Evolution, 2014).

33. C. J. Pennycuick, Gliding flight of the white-backed vulture Gyps africanus. J. Exp. Biol. 55, 13–38 (1971).

34. M. T. Wilkinson, Three-dimensional geometry of a pterosaur wing skeleton, and its implications for aerial and terrestrial locomotion. Zool. J. Linn. Soc. 154, 27–69 (2008).

35. S. C. Bennett, Morphological evolution of the pectoral girdle of pterosaurs: myology and function. Geol. Soc. London, Spec. Publ. 217, 191–215 (2003).

36. U. M. Norberg, J. M. V Rayner, Ecological morphology and flight in bats (Mammalia; Chiroptera): wing adaptations, flight performance, foraging strategy and echolocation. Philos. Trans. R. Soc. London. B, Biol. Sci. 316, 335–427 (1987).

37. J. M. V Rayner, in Current ornithology (Springer, 1988), pp. 1–66.

38. M. P. Witton, D. Naish, A reappraisal of azhdarchid pterosaur functional morphology and paleoecology. PLoS One. 3, e2271 (2008).

39. M. P. Witton, D. Naish, Azhdarchid pterosaurs: water-trawling pelican mimics or “terrestrial stalkers”? Acta Palaeontol. Pol. 60, 651–660 (2013).

40. H. Weimerskirch, C. Bishop, T. Jeanniard-du-Dot, A. Prudor, G. Sachs, Frigate birds track atmospheric conditions over months-long transoceanic flights. Science. 353, 74–78 (2016).

41. G. Sachs, H. Weimerskirch, Flight of frigatebirds inside clouds–energy gain, stability and control. J. Theor. Biol. 448, 9–16 (2018).

42. B. Swaminathan, R. Mohan, in 2018 Atmospheric Flight Mechanics Conference (2018), p. 2832.

43. M. P. Buckley, F. Veron, Structure of the airflow above surface waves. J. Phys. Oceanogr. 46, 1377–1397 (2016).

44. G. Sachs, J. Traugott, A. P. Nesterova, F. Bonadonna, Experimental verification of dynamic soaring in albatrosses. J. Exp. Biol. 216, 4222–4232 (2013).

45. R. Harel, N. Horvitz, R. Nathan, Adult vultures outperform juveniles in challenging thermal soaring conditions. Sci. Rep. 6, 1–8 (2016).

46. S. Sherub, G. Bohrer, M. Wikelski, R. Weinzierl, Behavioural adaptations to flight into thin air. Biol. Lett. 12, 20160432 (2016).

47. H. J. Williams, O. Duriez, M. D. Holton, G. Dell’Omo, R. P. Wilson, E. L. C. Shepard, Vultures respond to challenges of near-ground thermal soaring by varying bank angle. J. Exp. Biol. 221(2018).

48. I. K. Shimatani, K. Yoda, N. Katsumata, K. Sato, Toward the quantification of a conceptual framework for movement ecology using circular statistical modeling. PLoS One. 7, e50309 (2012).

49. J. Treep, G. Bohrer, J. Shamoun-Baranes, O. Duriez, R. Prata de Moraes Frasson, W. Bouten, Using high resolution GPS tracking data of bird flight for meteorological observations. Bull. Am. Meteorol. Soc. 97, 951–961 (2015).

50. R. Weinzierl, G. Bohrer, B. Kranstauber, W. Fiedler, M. Wikelski, A. Flack, Wind estimation based on thermal soaring of birds. Ecol. Evol. 6, 8706–8718 (2016).

51. Y. Yonehara, Y. Goto, K. Yoda, Y. Watanuki, L. C. Young, H. Weimerskirch, K. Sato, Flight paths of seabirds soaring over the ocean surface enable measurement of fine-scale wind speed and direction. Proc. Natl. Acad. Sci. U. S. A. 113, 9039–9044 (2016).

52. Y. Goto, K. Yoda, K. Sato, Asymmetry hidden in birds’ tracks reveals wind, heading, and orientation ability over the ocean. Sci. Adv. 3, e1700097 (2017).

53. C. Péron, K. Delord, R. A. Phillips, Y. Charbonnier, C. Marteau, M. Louzao, H. Weimerskirch, Seasonal variation in oceanographic habitat and behaviour of white-chinned petrels Procellaria aequinoctialis from Kerguelen Island. Mar. Ecol. Prog. Ser. 416, 267–284 (2010).

54. I. Mir, A. Maqsood, S. A. Eisa, H. Taha, S. Akhtar, Optimal morphing–augmented dynamic soaring maneuvers for unmanned air vehicle capable of span and sweep morphologies. Aerosp. Sci. Technol. 79, 17–36 (2018).

55. C. D. Bramwell, G. R. Whitfield, Biomechanics of pteranodon. Philos. Trans. R. Soc. London. B, Biol. Sci. 267, 503–581 (1974).

56. J. C. Brower, The aerodynamics of Pteranodon and Nyctosaurus, two large pterosaurs from the Upper Cretaceous of Kansas. J. Vertebr. Paleontol. 3, 84–124 (1983).

57. M. T. Wilkinson, D. M. Unwin, C. P. Ellington, High lift function of the pteroid bone and forewing of pterosaurs. Proc. R. Soc. B Biol. Sci. 273, 119–126 (2006).

58. C. Palmer, Inferring the properties of the pterosaur wing membrane. Geol. Soc. London, Spec. Publ. 455, 57–68 (2018).

59. J. J. Bird, J. W. Langelaan, C. Montella, J. Spletzer, J. L. Grenestedt, in AIAA Guidance, Navigation, and Control Conference (2014), p. 263.

60. J. Nocedal, S. Wright, Numerical optimization (Springer Science & Business Media, 2006).

61. U. M. Norberg, Vertebrate flight: mechanics, physiology, morphology, ecology and evolution (Springer Science & Business Media, 2012), vol. 27.

62. U. M. Lindhe-Norberg, A. P. Brooke, W. J. Trewhella, Soaring and non-soaring bats of the family Pteropodidae (flying foxes, Pteropus spp.): wing morphology and flight performance. J. Exp. Biol. 203, 651–664 (2000).

63. S. A. Shaffer, H. Weimerskirch, D. P. Costa, Functional significance of sexual dimorphism in wandering albatrosses, Diomedea exulans. Funct. Ecol. 15, 203–210 (2001).

64. R. A. Phillips, J. R. D. Silk, B. Phalan, P. Catry, J. P. Croxall, Seasonal sexual segregation in two Thalassarche albatross species: competitive exclusion, reproductive role specialization or foraging niche divergence? Proc. R. Soc. London. Ser. B Biol. Sci. 271, 1283–1291 (2004).

65. C. J. Pennycuick, Flight of auks (Alcidae) and other northern seabirds compared with southern Procellariiformes: ornithodolite observations. J. Exp. Biol. 128, 335–347 (1987).

