## Supplementary Information for "Soaring styles of extinct giant birds and pterosaurs"

### Supplementary Materials

Fig. S1. Glide polars and circling envelopes where $C_L^*$ is the lift coefficient at the minimum sinking speed assuming a wing-drag minimizing wingspan reduction.

Fig. S2. Glide polars and circling envelopes where $C_L^*$ is the lift coefficient at the minimum sinking speed assuming a fixed wingspan.

Fig. S3. Glide polars and circling envelopes where $C_L^*$ is the maximum lift coefficient assuming a linear wingspan reduction.

Fig. S4. Glide polars and circling envelopes where $C_L^*$ is at the maximum lift coefficient.

Fig. S5. Glide polars and circling envelopes where $C_L^*$ is at the maximum lift coefficient.

Table S1. Thermal soaring performances where $C_L^*$ is at the minimum sinking speed assuming linear wingspan reduction

Table S2. Thermal soaring performances where $C_L^*$ is at the minimum sinking speed assuming a wing-drag minimizing wingspan.

Table S3. Thermal soaring performances where $C_L^*$ is at the minimum sinking speed assuming a fixed wingspan.

Table S4. Thermal soaring performances where $C_L^*$ is the maximum lift coefficient assuming linear wingspan reduction.

Table S5. Thermal soaring performances where $C_L^*$ is the maximum lift coefficient assuming a wing-drag minimizing wingspan.

Table S6. Thermal soaring performances where $C_L^*$ is the maximum lift coefficient assuming a fixed wingspan.

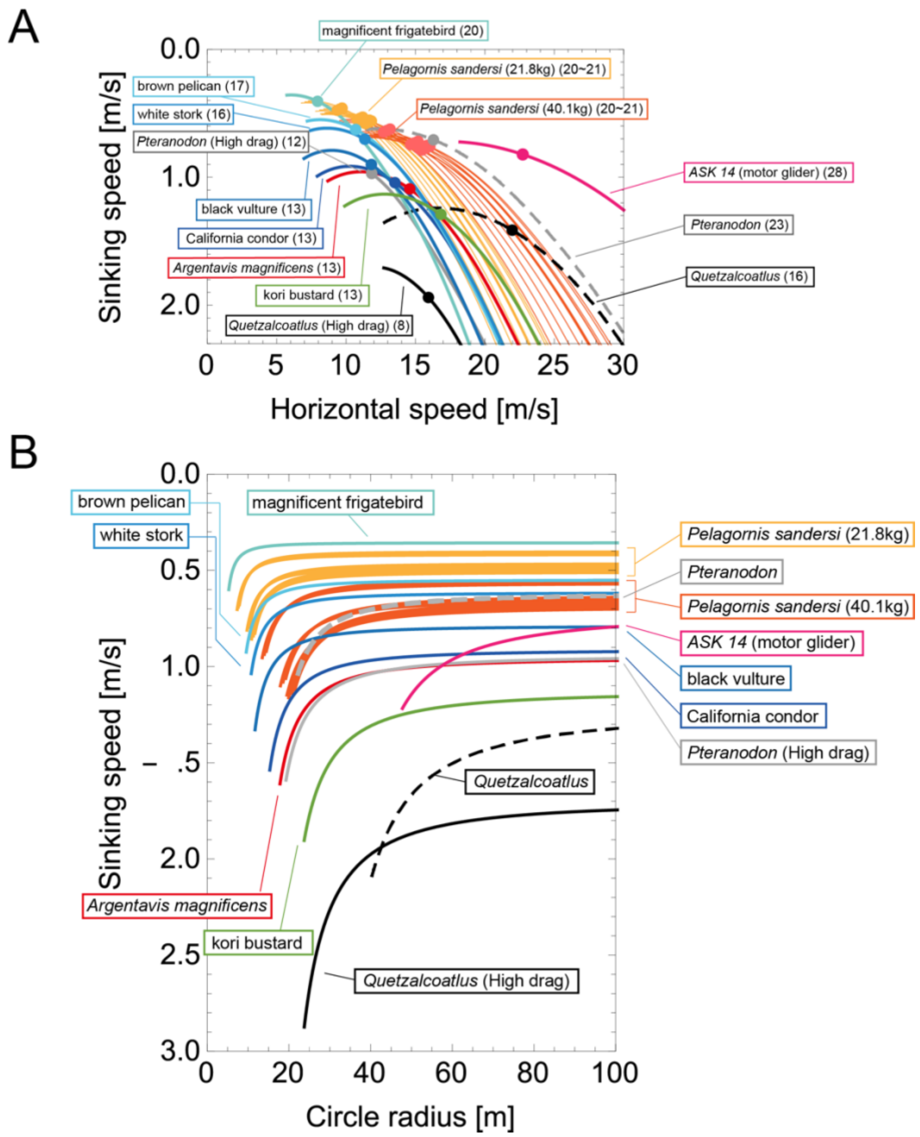


**Fig. S1. Glide polars and circling envelopes where $C_L^*$ is the lift coefficient at the minimum sinking speed assuming a wing-drag minimizing wingspan reduction.** (A) Glide polars and (B) circling envelopes of extinct species, extant thermal soaring species, and the kori bustard, the heaviest rarely flying bird, assuming a wing-drag minimizing wingspan reduction ($\beta = \min\left(1, 2\left[\frac{km^2g^2}{\pi\rho^2b^2C_{\mathrm{Dpro}}S_W V^4}\right]^{\frac{1}{3}}\right)$). In (A), the maximum glide ratios of each species are shown on the right side of species names. Points represent the horizontal speed and sinking speed at the maximum glide ratio of each species. (B) shows a circling envelope with a bank angle of up to 45°. The circle radius becomes smaller as the bank angle is increased. The lift coefficient of circling envelope ( $C_L^*$) is the lift coefficient at the minimum sinking speed.

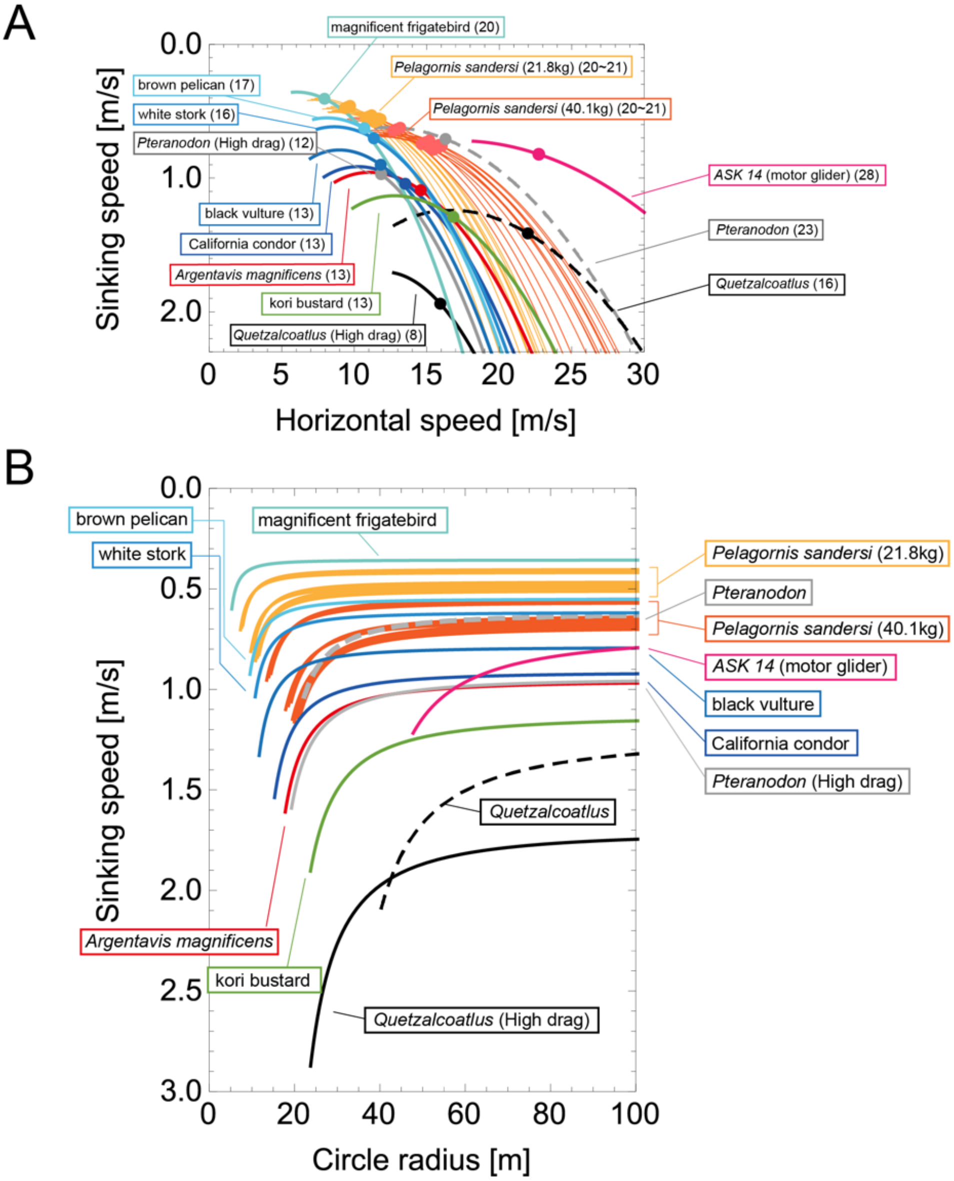


**Fig. S2. Glide polars and circling envelopes where $C_L^*$ is the lift coefficient at the minimum sinking speed assuming a fixed wingspan.** (A) Glide polars and (B) circling envelopes of extinct species, extant thermal soaring species, and the kori bustard, the heaviest rarely flying bird, assuming a fixed wingspan ($\beta = 1$). In (A), the maximum glide ratios of each species are shown on the right side of species names. Points represent the horizontal speed and sinking speed at the maximum glide ratio of each species. (B) shows a circling envelope with a bank angle of up to 45°. The circle radius becomes smaller as the bank angle is increased. The lift coefficient of circling envelope ( $C_L^*$) is the lift coefficient at the minimum sinking speed.

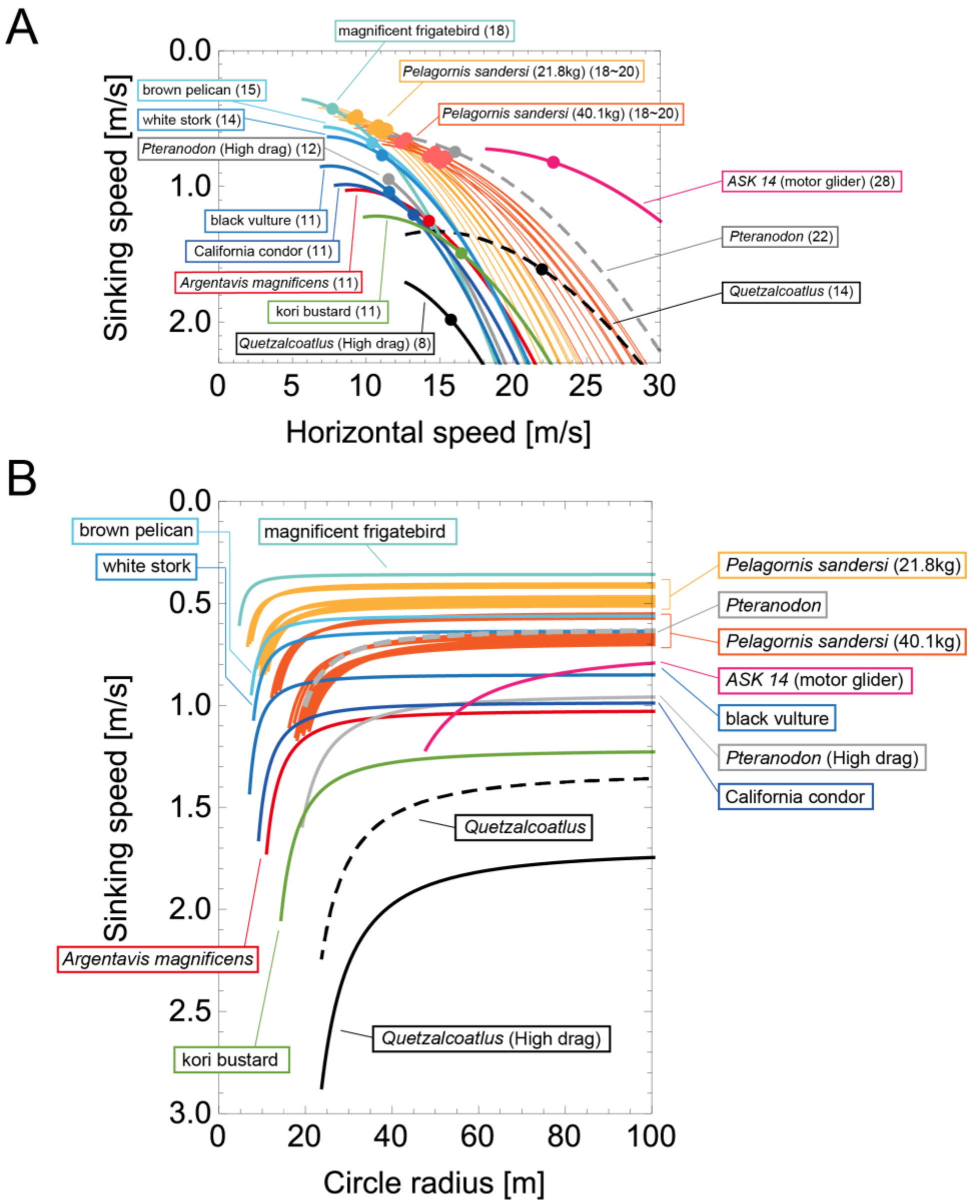


**Fig. S3. Glide polars and circling envelopes where $C_L^*$ is the maximum lift coefficient assuming a linear wingspan reduction.** (A) Glide polars and (B) circling envelopes of extinct species, extant thermal soaring species, and the kori bustard, the heaviest rarely flying bird, assuming a linear wingspan reduction. In (A), the maximum glide ratios of each species are shown on the right side of species names. Points represent the horizontal speed and sinking speed at the maximum glide ratio of each species. (B) shows a circling envelope with a bank angle of up to 45°. The circle radius becomes smaller as the bank angle is increased. The lift coefficient of circling envelope ( $C_L^*$) is the maximum lift coefficient.

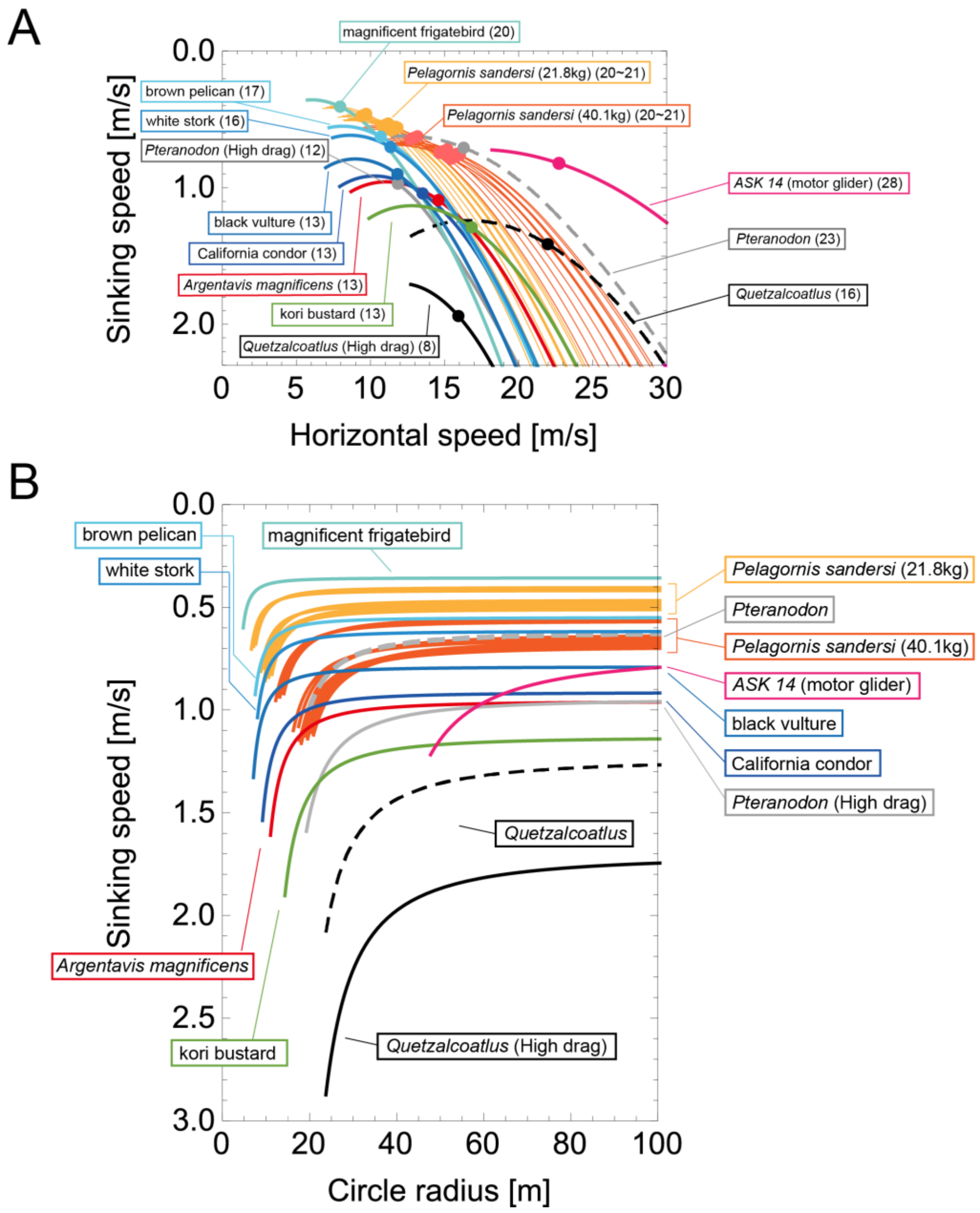


**Fig. S4. Glide polars and circling envelopes where $C_L^*$ is the maximum lift coefficient assuming a wing-drag minimizing wingspan reduction.** (A) Glide polars and (B) circling envelopes of extinct species, extant thermal soaring species, and the kori bustard, the heaviest rarely flying bird, assuming a wing-drag minimizing wingspan reduction). In (A), the maximum glide ratios of each species are shown on the right side of species names. Points represent the horizontal speed and sinking speed at the maximum glide ratio of each species. (B) shows a circling envelope with a bank angle of up to 45°. The circle radius becomes smaller as the bank angle is increased. The lift coefficient of circling envelope ( $C_L^*$) is the maximum lift coefficient.

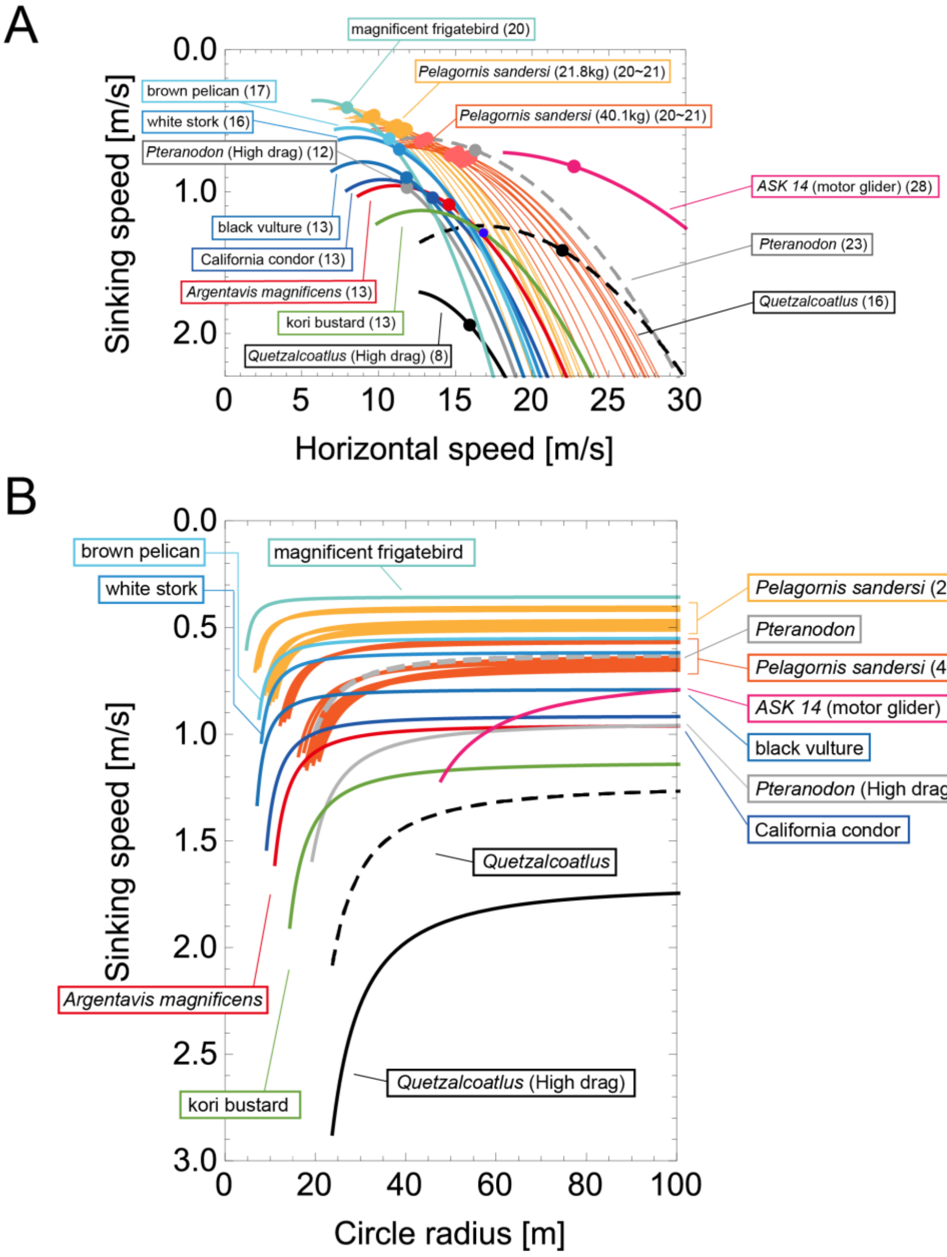


**Fig. S5. Glide polars and circling envelopes where $C_L^*$ is the maximum lift coefficient assuming a fixed wingspan.** (A) Glide polars and (B) circling envelopes of extinct species, extant thermal soaring species, and the kori bustard, the heaviest rarely flying bird, assuming a fixed wingspan ($\beta = 1$). In (A), the maximum glide ratios of each species are shown on the right side of species names. Points represent the horizontal speed and sinking speed at the maximum glide ratio of each species. (B) Shows a circling envelope with a bank angle of up to 45°. The circle radius becomes smaller as the bank angle is increased. The lift coefficient of circling envelope ( $C_L^*$) is the maximum lift coefficient.

| Species | Maximum glide ratio | Horizontal speed at maximum glide ratio [m/s] | Limitting radius [m] | Minimum sinnking speed [m/s] | Lift coefficient of circling envelope $C_L$* |
|---|---|---|---|---|---|
| Pelagornis sandersi (Mass 21.8 kg Wingspan 6.06 m Aspect ratio 13) | 18.1 | 11.1 | 6.96 | 0.511 | 1.80 |
| Pelagornis sandersi (Mass 21.8 kg Wingspan 6.06 m Aspect ratio 14) | 19.0 | 11.3 | 7.52 | 0.501 | 1.80 |
| Pelagornis sandersi (Mass 21.8 kg Wingspan 6.06 m Aspect ratio 15) | 19.7 | 11.4 | 8.04 | 0.492 | 1.80 |
| Pelagornis sandersi (Mass 21.8 kg Wingspan 6.13 m Aspect ratio 13) | 18.2 | 10.9 | 6.81 | 0.505 | 1.80 |
| Pelagornis sandersi (Mass 21.8 kg Wingspan 6.13 m Aspect ratio 14) | 19.0 | 11.1 | 7.35 | 0.495 | 1.80 |
| Pelagornis sandersi (Mass 21.8 kg Wingspan 6.13 m Aspect ratio 15) | 19.7 | 11.3 | 7.85 | 0.486 | 1.80 |
| Pelagornis sandersi (Mass 21.8 kg Wingspan 6.40 m Aspect ratio 13) | 18.2 | 10.5 | 6.25 | 0.484 | 1.80 |
| Pelagornis sandersi (Mass 21.8 kg Wingspan 6.40 m Aspect ratio 14) | 18.9 | 10.7 | 6.72 | 0.474 | 1.80 |
| Pelagornis sandersi (Mass 21.8 kg Wingspan 6.40 m Aspect ratio 15) | 19.7 | 10.8 | 7.21 | 0.466 | 1.80 |
| Pelagornis sandersi (Mass 21.8 kg Wingspan 7.38 m Aspect ratio 13) | 18.2 | 9.09 | 4.70 | 0.420 | 1.80 |
| Pelagornis sandersi (Mass 21.8 kg Wingspan 7.38 m Aspect ratio 14) | 19.0 | 9.25 | 5.06 | 0.411 | 1.80 |
| Pelagornis sandersi (Mass 21.8 kg Wingspan 7.38 m Aspect ratio 15) | 19.7 | 9.40 | 5.43 | 0.404 | 1.80 |
| Pelagornis sandersi (Mass 40.1 kg Wingspan 6.06 m Aspect ratio 13) | 18.1 | 15.0 | 12.8 | 0.694 | 1.80 |
| Pelagornis sandersi (Mass 40.1 kg Wingspan 6.06 m Aspect ratio 14) | 19.0 | 15.3 | 13.8 | 0.679 | 1.80 |
| Pelagornis sandersi (Mass 40.1 kg Wingspan 6.06 m Aspect ratio 15) | 19.7 | 15.5 | 14.8 | 0.667 | 1.80 |
| Pelagornis sandersi (Mass 40.1 kg Wingspan 6.13 m Aspect ratio 13) | 18.2 | 14.8 | 12.5 | 0.685 | 1.80 |
| Pelagornis sandersi (Mass 40.1 kg Wingspan 6.13 m Aspect ratio 14) | 19.0 | 15.1 | 13.5 | 0.671 | 1.80 |
| Pelagornis sandersi (Mass 40.1 kg Wingspan 6.13 m Aspect ratio 15) | 19.7 | 15.3 | 14.4 | 0.660 | 1.80 |
| Pelagornis sandersi (Mass 40.1 kg Wingspan 6.40 m Aspect ratio 13) | 18.2 | 14.2 | 11.5 | 0.656 | 1.80 |
| Pelagornis sandersi (Mass 40.1 kg Wingspan 6.40 m Aspect ratio 14) | 18.9 | 14.5 | 12.4 | 0.643 | 1.80 |
| Pelagornis sandersi (Mass 40.1 kg Wingspan 6.40 m Aspect ratio 15) | 19.7 | 14.7 | 13.3 | 0.631 | 1.80 |
| Pelagornis sandersi (Mass 40.1 kg Wingspan 7.38 m Aspect ratio 13) | 18.2 | 12.3 | 8.65 | 0.569 | 1.80 |
| Pelagornis sandersi (Mass 40.1 kg Wingspan 7.38 m Aspect ratio 14) | 19.0 | 12.5 | 9.31 | 0.558 | 1.80 |
| Pelagornis sandersi (Mass 40.1 kg Wingspan 7.38 m Aspect ratio 15) | 19.7 | 12.7 | 9.98 | 0.548 | 1.80 |
| *Argentavis magnificens* | 11.4 | 14.3 | 8.77 | 1.02 | 1.60 |
| *Pteranodon* | 21.5 | 16.0 | 13.6 | 0.625 | 2.20 |
| *Pteranodon* (high drag) | 12.2 | 11.5 | 13.6 | 0.946 | 2.20 |
| *Quetzalcoatlus* | 13.6 | 22.0 | 21.4 | 1.33 | 1.73 |
| *Quetzalcoatlus* (high drag) | 7.96 | 15.8 | 16.8 | 1.71 | 2.20 |
| magnificent frigatebird | 18.0 | 7.71 | 3.37 | 0.358 | 1.80 |
| California condor | 11.0 | 13.2 | 7.48 | 0.986 | 1.57 |
| brown pelican | 15.3 | 10.4 | 5.32 | 0.561 | 1.80 |
| black vulture | 11.1 | 11.6 | 5.73 | 0.849 | 1.58 |
| white strok | 14.4 | 11.1 | 5.69 | 0.635 | 1.80 |
| kori bustard | 11.0 | 16.5 | 11.6 | 1.22 | 1.57 |

**Table S1. Thermal soaring performances where $C_L^*$ is the lift coefficient at the minimum sinking speed assuming linear wingspan reduction.** Maximum glide ratio, and horizontal speed at maximum glide ratio limiting radius, minimum sinking speed, and lift coefficients of circling envelopes assuming linear wingspan reduction.

| Species | Maximum glide ratio | Horizontal speed at maximum glide ratio [m/s] | Limiting radius [m] | Minimum sinking speed [m/s] | Lift coefficient of circling envelope $C_L$* |
|---|---|---|---|---|---|
| Pelagornis sandersi (Mass 21.8 kg Wingspan 6.06 m Aspect ratio 13) | 19.6 | 11.4 | 7.67 | 0.510 | 1.63 |
| Pelagornis sandersi (Mass 21.8 kg Wingspan 6.06 m Aspect ratio 14) | 20.4 | 11.6 | 7.97 | 0.500 | 1.70 |
| Pelagornis sandersi (Mass 21.8 kg Wingspan 6.06 m Aspect ratio 15) | 21.1 | 11.8 | 8.24 | 0.492 | 1.76 |
| Pelagornis sandersi (Mass 21.8 kg Wingspan 6.13 m Aspect ratio 13) | 19.6 | 11.3 | 7.50 | 0.504 | 1.64 |
| Pelagornis sandersi (Mass 21.8 kg Wingspan 6.13 m Aspect ratio 14) | 20.4 | 11.5 | 7.79 | 0.494 | 1.70 |
| Pelagornis sandersi (Mass 21.8 kg Wingspan 6.13 m Aspect ratio 15) | 21.1 | 11.7 | 8.05 | 0.486 | 1.75 |
| Pelagornis sandersi (Mass 21.8 kg Wingspan 6.40 m Aspect ratio 13) | 19.6 | 10.8 | 6.88 | 0.482 | 1.64 |
| Pelagornis sandersi (Mass 21.8 kg Wingspan 6.40 m Aspect ratio 14) | 20.4 | 11.0 | 7.14 | 0.474 | 1.70 |
| Pelagornis sandersi (Mass 21.8 kg Wingspan 6.40 m Aspect ratio 15) | 21.1 | 11.2 | 7.39 | 0.465 | 1.76 |
| Pelagornis sandersi (Mass 21.8 kg Wingspan 7.38 m Aspect ratio 13) | 19.6 | 9.36 | 5.17 | 0.418 | 1.63 |
| Pelagornis sandersi (Mass 21.8 kg Wingspan 7.38 m Aspect ratio 14) | 20.4 | 9.54 | 5.37 | 0.411 | 1.70 |
| Pelagornis sandersi (Mass 21.8 kg Wingspan 7.38 m Aspect ratio 15) | 21.1 | 9.71 | 5.56 | 0.404 | 1.76 |
| Pelagornis sandersi (Mass 40.1 kg Wingspan 6.06 m Aspect ratio 13) | 19.6 | 15.5 | 14.1 | 0.691 | 1.63 |
| Pelagornis sandersi (Mass 40.1 kg Wingspan 6.06 m Aspect ratio 14) | 20.4 | 15.8 | 14.7 | 0.678 | 1.70 |
| Pelagornis sandersi (Mass 40.1 kg Wingspan 6.06 m Aspect ratio 15) | 21.1 | 16.0 | 15.2 | 0.667 | 1.76 |
| Pelagornis sandersi (Mass 40.1 kg Wingspan 6.13 m Aspect ratio 13) | 19.6 | 15.3 | 13.8 | 0.683 | 1.64 |
| Pelagornis sandersi (Mass 40.1 kg Wingspan 6.13 m Aspect ratio 14) | 20.4 | 15.6 | 14.3 | 0.670 | 1.70 |
| Pelagornis sandersi (Mass 40.1 kg Wingspan 6.13 m Aspect ratio 15) | 21.1 | 15.8 | 14.8 | 0.659 | 1.75 |
| Pelagornis sandersi (Mass 40.1 kg Wingspan 6.40 m Aspect ratio 13) | 19.6 | 14.6 | 12.7 | 0.654 | 1.64 |
| Pelagornis sandersi (Mass 40.1 kg Wingspan 6.40 m Aspect ratio 14) | 20.4 | 14.9 | 13.1 | 0.642 | 1.70 |
| Pelagornis sandersi (Mass 40.1 kg Wingspan 6.40 m Aspect ratio 15) | 21.1 | 15.2 | 13.6 | 0.631 | 1.76 |
| Pelagornis sandersi (Mass 40.1 kg Wingspan 7.38 m Aspect ratio 13) | 19.6 | 12.7 | 9.52 | 0.567 | 1.63 |
| Pelagornis sandersi (Mass 40.1 kg Wingspan 7.38 m Aspect ratio 14) | 20.4 | 12.9 | 9.88 | 0.557 | 1.70 |
| Pelagornis sandersi (Mass 40.1 kg Wingspan 7.38 m Aspect ratio 15) | 21.1 | 13.2 | 10.2 | 0.547 | 1.76 |
| *Argentavis magnificens* | 13.4 | 14.6 | 12.6 | 0.957 | 1.11 |
| *Pteranodon* | 23.0 | 16.3 | 15.7 | 0.621 | 1.92 |
| *Pteranodon* (high drag) | 12.2 | 11.8 | 13.6 | 0.946 | 2.20 |
| *Quetzalcoatlus* | 15.5 | 22.0 | 28.5 | 1.24 | 1.29 |
| *Quetzalcoatlus* (high drag) | 8.21 | 15.9 | 16.8 | 1.71 | 2.20 |
| magnificent frigatebird | 19.5 | 7.95 | 3.73 | 0.357 | 1.63 |
| California condor | 13.0 | 13.5 | 10.8 | 0.915 | 1.08 |
| brown pelican | 17.0 | 10.7 | 6.75 | 0.550 | 1.42 |
| black vulture | 13.1 | 11.8 | 8.27 | 0.790 | 1.09 |
| white strok | 16.1 | 11.3 | 7.61 | 0.617 | 1.35 |
| kori bustard | 13.0 | 16.8 | 16.8 | 1.13 | 1.09 |

**Table S2. Thermal soaring performances where $C_L^*$ is the lift coefficient at the minimum sinking speed assuming a wing-drag minimizing wingspan.** Maximum glide ratio, and horizontal speed at maximum glide ratio limiting radius, minimum sinking speed, and lift coefficients of circling envelopes assuming a wing-drag minimizing wingspan.

| Species | Maximum glide ratio | Horizontal speed at maximum glide ratio [m/s] | Limiting radius [m] | Minimum sinking speed [m/s] | Lift coefficient of circling envelope $C_L$* |
|---|---|---|---|---|---|
| Pelagornis sandersi (Mass 21.8 kg Wingspan 6.06 m Aspect ratio 13) | 19.6 | 11.4 | 7.67 | 0.510 | 1.63 |
| Pelagornis sandersi (Mass 21.8 kg Wingspan 6.06 m Aspect ratio 14) | 20.4 | 11.6 | 7.97 | 0.500 | 1.70 |
| Pelagornis sandersi (Mass 21.8 kg Wingspan 6.06 m Aspect ratio 15) | 21.1 | 11.8 | 8.24 | 0.492 | 1.76 |
| Pelagornis sandersi (Mass 21.8 kg Wingspan 6.13 m Aspect ratio 13) | 19.6 | 11.3 | 7.50 | 0.504 | 1.64 |
| Pelagornis sandersi (Mass 21.8 kg Wingspan 6.13 m Aspect ratio 14) | 20.4 | 11.5 | 7.79 | 0.494 | 1.70 |
| Pelagornis sandersi (Mass 21.8 kg Wingspan 6.13 m Aspect ratio 15) | 21.1 | 11.7 | 8.05 | 0.486 | 1.75 |
| Pelagornis sandersi (Mass 21.8 kg Wingspan 6.40 m Aspect ratio 13) | 19.6 | 10.8 | 6.88 | 0.482 | 1.64 |
| Pelagornis sandersi (Mass 21.8 kg Wingspan 6.40 m Aspect ratio 14) | 20.4 | 11.0 | 7.14 | 0.474 | 1.70 |
| Pelagornis sandersi (Mass 21.8 kg Wingspan 6.40 m Aspect ratio 15) | 21.1 | 11.2 | 7.39 | 0.465 | 1.76 |
| Pelagornis sandersi (Mass 21.8 kg Wingspan 7.38 m Aspect ratio 13) | 19.6 | 9.36 | 5.17 | 0.418 | 1.63 |
| Pelagornis sandersi (Mass 21.8 kg Wingspan 7.38 m Aspect ratio 14) | 20.4 | 9.54 | 5.37 | 0.411 | 1.70 |
| Pelagornis sandersi (Mass 21.8 kg Wingspan 7.38 m Aspect ratio 15) | 21.1 | 9.71 | 5.56 | 0.404 | 1.76 |
| Pelagornis sandersi (Mass 40.1 kg Wingspan 6.06 m Aspect ratio 13) | 19.6 | 15.5 | 14.1 | 0.691 | 1.63 |
| Pelagornis sandersi (Mass 40.1 kg Wingspan 6.06 m Aspect ratio 14) | 20.4 | 15.8 | 14.7 | 0.678 | 1.70 |
| Pelagornis sandersi (Mass 40.1 kg Wingspan 6.06 m Aspect ratio 15) | 21.1 | 16.0 | 15.2 | 0.667 | 1.76 |
| Pelagornis sandersi (Mass 40.1 kg Wingspan 6.13 m Aspect ratio 13) | 19.6 | 15.3 | 13.8 | 0.683 | 1.64 |
| Pelagornis sandersi (Mass 40.1 kg Wingspan 6.13 m Aspect ratio 14) | 20.4 | 15.6 | 14.3 | 0.670 | 1.70 |
| Pelagornis sandersi (Mass 40.1 kg Wingspan 6.13 m Aspect ratio 15) | 21.1 | 15.8 | 14.8 | 0.659 | 1.75 |
| Pelagornis sandersi (Mass 40.1 kg Wingspan 6.40 m Aspect ratio 13) | 19.6 | 14.6 | 12.7 | 0.654 | 1.64 |
| Pelagornis sandersi (Mass 40.1 kg Wingspan 6.40 m Aspect ratio 14) | 20.4 | 14.9 | 13.1 | 0.642 | 1.70 |
| Pelagornis sandersi (Mass 40.1 kg Wingspan 6.40 m Aspect ratio 15) | 21.1 | 15.2 | 13.6 | 0.631 | 1.76 |
| Pelagornis sandersi (Mass 40.1 kg Wingspan 7.38 m Aspect ratio 13) | 19.6 | 12.7 | 9.52 | 0.567 | 1.63 |
| Pelagornis sandersi (Mass 40.1 kg Wingspan 7.38 m Aspect ratio 14) | 20.4 | 12.9 | 9.88 | 0.557 | 1.70 |
| Pelagornis sandersi (Mass 40.1 kg Wingspan 7.38 m Aspect ratio 15) | 21.1 | 13.2 | 10.2 | 0.547 | 1.76 |
| *Argentavis magnificens* | 13.4 | 14.6 | 12.6 | 0.957 | 1.11 |
| *Pteranodon* | 23.0 | 16.3 | 15.7 | 0.621 | 1.92 |
| *Pteranodon* (high drag) | 12.2 | 11.9 | 13.6 | 0.946 | 2.20 |
| *Quetzalcoatlus* | 15.5 | 22.0 | 28.5 | 1.24 | 1.29 |
| *Quetzalcoatlus* (high drag) | 8.21 | 15.9 | 16.8 | 1.71 | 2.20 |
| magnificent frigatebird | 19.5 | 7.95 | 3.73 | 0.357 | 1.63 |
| California condor | 13.0 | 13.5 | 10.8 | 0.915 | 1.08 |
| brown pelican | 17.0 | 10.7 | 6.75 | 0.550 | 1.42 |
| black vulture | 13.1 | 11.8 | 8.27 | 0.790 | 1.09 |
| white strok | 16.1 | 11.3 | 7.61 | 0.617 | 1.35 |
| kori bustard | 13.0 | 16.8 | 16.8 | 1.13 | 1.09 |

**Table S3. Thermal soaring performances where $C_L^*$ is the lift coefficient at the minimum sinking speed assuming a fixed wingspan.** Maximum glide ratio, and horizontal speed at maximum glide ratio limiting radius, minimum sinking speed, and lift coefficients of circling envelopes assuming a fixed wingspan.

| Species | Maximum glide ratio | Horizontal speed at maximum glide ratio [m/s] | Limiting radius [m] | Minimum sinking speed [m/s] | Lift coefficient of circling envelope $C_L$* |
|---|---|---|---|---|---|
| Pelagornis sandersi (Mass 21.8 kg Wingspan 6.06 m Aspect ratio 13) | 18.1 | 11.1 | 6.96 | 0.511 | 1.80 |
| Pelagornis sandersi (Mass 21.8 kg Wingspan 6.06 m Aspect ratio 14) | 19.0 | 11.3 | 7.52 | 0.501 | 1.80 |
| Pelagornis sandersi (Mass 21.8 kg Wingspan 6.06 m Aspect ratio 15) | 19.7 | 11.4 | 8.04 | 0.492 | 1.80 |
| Pelagornis sandersi (Mass 21.8 kg Wingspan 6.13 m Aspect ratio 13) | 18.2 | 10.9 | 6.81 | 0.505 | 1.80 |
| Pelagornis sandersi (Mass 21.8 kg Wingspan 6.13 m Aspect ratio 14) | 19.0 | 11.1 | 7.35 | 0.495 | 1.80 |
| Pelagornis sandersi (Mass 21.8 kg Wingspan 6.13 m Aspect ratio 15) | 19.7 | 11.3 | 7.85 | 0.486 | 1.80 |
| Pelagornis sandersi (Mass 21.8 kg Wingspan 6.40 m Aspect ratio 13) | 18.2 | 10.5 | 6.25 | 0.484 | 1.80 |
| Pelagornis sandersi (Mass 21.8 kg Wingspan 6.40 m Aspect ratio 14) | 18.9 | 10.7 | 6.72 | 0.474 | 1.80 |
| Pelagornis sandersi (Mass 21.8 kg Wingspan 6.40 m Aspect ratio 15) | 19.7 | 10.8 | 7.21 | 0.466 | 1.80 |
| Pelagornis sandersi (Mass 21.8 kg Wingspan 7.38 m Aspect ratio 13) | 18.2 | 9.09 | 4.70 | 0.420 | 1.80 |
| Pelagornis sandersi (Mass 21.8 kg Wingspan 7.38 m Aspect ratio 14) | 19.0 | 9.25 | 5.06 | 0.411 | 1.80 |
| Pelagornis sandersi (Mass 21.8 kg Wingspan 7.38 m Aspect ratio 15) | 19.7 | 9.40 | 5.43 | 0.404 | 1.80 |
| Pelagornis sandersi (Mass 40.1 kg Wingspan 6.06 m Aspect ratio 13) | 18.1 | 15.0 | 12.8 | 0.694 | 1.80 |
| Pelagornis sandersi (Mass 40.1 kg Wingspan 6.06 m Aspect ratio 14) | 19.0 | 15.3 | 13.8 | 0.679 | 1.80 |
| Pelagornis sandersi (Mass 40.1 kg Wingspan 6.06 m Aspect ratio 15) | 19.7 | 15.5 | 14.8 | 0.667 | 1.80 |
| Pelagornis sandersi (Mass 40.1 kg Wingspan 6.13 m Aspect ratio 13) | 18.2 | 14.8 | 12.5 | 0.685 | 1.80 |
| Pelagornis sandersi (Mass 40.1 kg Wingspan 6.13 m Aspect ratio 14) | 19.0 | 15.1 | 13.5 | 0.671 | 1.80 |
| Pelagornis sandersi (Mass 40.1 kg Wingspan 6.13 m Aspect ratio 15) | 19.7 | 15.3 | 14.4 | 0.660 | 1.80 |
| Pelagornis sandersi (Mass 40.1 kg Wingspan 6.40 m Aspect ratio 13) | 18.2 | 14.2 | 11.5 | 0.656 | 1.80 |
| Pelagornis sandersi (Mass 40.1 kg Wingspan 6.40 m Aspect ratio 14) | 18.9 | 14.5 | 12.4 | 0.643 | 1.80 |
| Pelagornis sandersi (Mass 40.1 kg Wingspan 6.40 m Aspect ratio 15) | 19.7 | 14.7 | 13.3 | 0.631 | 1.80 |
| Pelagornis sandersi (Mass 40.1 kg Wingspan 7.38 m Aspect ratio 13) | 18.2 | 12.3 | 8.65 | 0.569 | 1.80 |
| Pelagornis sandersi (Mass 40.1 kg Wingspan 7.38 m Aspect ratio 14) | 19.0 | 12.5 | 9.31 | 0.558 | 1.80 |
| Pelagornis sandersi (Mass 40.1 kg Wingspan 7.38 m Aspect ratio 15) | 19.7 | 12.7 | 10.0 | 0.548 | 1.80 |
| *Argentavis magnificens* | 11.4 | 14.3 | 7.80 | 1.02 | 1.80 |
| *Pteranodon* | 21.5 | 16.0 | 13.6 | 0.625 | 2.20 |
| *Pteranodon* (high drag) | 12.2 | 11.5 | 13.6 | 0.946 | 2.20 |
| *Quetzalcoatlus* | 13.6 | 22.0 | 16.8 | 1.33 | 2.20 |
| *Quetzalcoatlus* (high drag) | 7.96 | 15.8 | 16.8 | 1.71 | 2.20 |
| magnificent frigatebird | 18.0 | 7.71 | 3.37 | 0.358 | 1.80 |
| California condor | 11.0 | 13.2 | 6.50 | 0.986 | 1.80 |
| brown pelican | 15.3 | 10.4 | 5.32 | 0.561 | 1.80 |
| black vulture | 11.1 | 11.6 | 5.03 | 0.849 | 1.80 |
| white strok | 14.4 | 11.1 | 5.69 | 0.635 | 1.80 |
| kori bustard | 11.0 | 16.5 | 10.1 | 1.22 | 1.80 |

**Table S4. Thermal soaring performances where $C_L^*$ is the maximum lift coefficient assuming linear wingspan reduction.** Maximum glide ratio, and horizontal speed at maximum glide ratio limiting radius, minimum sinking speed, and lift coefficients of circling envelopes assuming linear wingspan reduction.

| Species | Maximum glide ratio | Horizontal speed at maximum glide ratio [m/s] | Limiting radius [m] | Minimum sinking speed [m/s] | Lift coefficient of circling envelope $C_L$* |
|---|---|---|---|---|---|
| Pelagornis sandersi (Mass 21.8 kg Wingspan 6.06 m Aspect ratio 13) | 19.6 | 11.4 | 6.96 | 0.510 | 1.80 |
| Pelagornis sandersi (Mass 21.8 kg Wingspan 6.06 m Aspect ratio 14) | 20.4 | 11.6 | 7.52 | 0.500 | 1.80 |
| Pelagornis sandersi (Mass 21.8 kg Wingspan 6.06 m Aspect ratio 15) | 21.1 | 11.8 | 8.04 | 0.492 | 1.80 |
| Pelagornis sandersi (Mass 21.8 kg Wingspan 6.13 m Aspect ratio 13) | 19.6 | 11.3 | 6.81 | 0.504 | 1.80 |
| Pelagornis sandersi (Mass 21.8 kg Wingspan 6.13 m Aspect ratio 14) | 20.4 | 11.5 | 7.35 | 0.494 | 1.80 |
| Pelagornis sandersi (Mass 21.8 kg Wingspan 6.13 m Aspect ratio 15) | 21.1 | 11.7 | 7.85 | 0.486 | 1.80 |
| Pelagornis sandersi (Mass 21.8 kg Wingspan 6.40 m Aspect ratio 13) | 19.6 | 10.8 | 6.25 | 0.482 | 1.80 |
| Pelagornis sandersi (Mass 21.8 kg Wingspan 6.40 m Aspect ratio 14) | 20.4 | 11.0 | 6.72 | 0.474 | 1.80 |
| Pelagornis sandersi (Mass 21.8 kg Wingspan 6.40 m Aspect ratio 15) | 21.1 | 11.2 | 7.21 | 0.465 | 1.80 |
| Pelagornis sandersi (Mass 21.8 kg Wingspan 7.38 m Aspect ratio 13) | 19.6 | 9.36 | 4.70 | 0.418 | 1.80 |
| Pelagornis sandersi (Mass 21.8 kg Wingspan 7.38 m Aspect ratio 14) | 20.4 | 9.54 | 5.06 | 0.411 | 1.80 |
| Pelagornis sandersi (Mass 21.8 kg Wingspan 7.38 m Aspect ratio 15) | 21.1 | 9.71 | 5.43 | 0.404 | 1.80 |
| Pelagornis sandersi (Mass 40.1 kg Wingspan 6.06 m Aspect ratio 13) | 19.6 | 15.5 | 12.8 | 0.691 | 1.80 |
| Pelagornis sandersi (Mass 40.1 kg Wingspan 6.06 m Aspect ratio 14) | 20.4 | 15.8 | 13.8 | 0.678 | 1.80 |
| Pelagornis sandersi (Mass 40.1 kg Wingspan 6.06 m Aspect ratio 15) | 21.1 | 16.0 | 14.8 | 0.667 | 1.80 |
| Pelagornis sandersi (Mass 40.1 kg Wingspan 6.13 m Aspect ratio 13) | 19.6 | 15.3 | 12.5 | 0.683 | 1.80 |
| Pelagornis sandersi (Mass 40.1 kg Wingspan 6.13 m Aspect ratio 14) | 20.4 | 15.6 | 13.5 | 0.670 | 1.80 |
| Pelagornis sandersi (Mass 40.1 kg Wingspan 6.13 m Aspect ratio 15) | 21.1 | 15.8 | 14.4 | 0.659 | 1.80 |
| Pelagornis sandersi (Mass 40.1 kg Wingspan 6.40 m Aspect ratio 13) | 19.6 | 14.6 | 11.5 | 0.654 | 1.80 |
| Pelagornis sandersi (Mass 40.1 kg Wingspan 6.40 m Aspect ratio 14) | 20.4 | 14.9 | 12.4 | 0.642 | 1.80 |
| Pelagornis sandersi (Mass 40.1 kg Wingspan 6.40 m Aspect ratio 15) | 21.1 | 15.2 | 13.3 | 0.631 | 1.80 |
| Pelagornis sandersi (Mass 40.1 kg Wingspan 7.38 m Aspect ratio 13) | 19.6 | 12.7 | 8.65 | 0.567 | 1.80 |
| Pelagornis sandersi (Mass 40.1 kg Wingspan 7.38 m Aspect ratio 14) | 20.4 | 12.9 | 9.31 | 0.557 | 1.80 |
| Pelagornis sandersi (Mass 40.1 kg Wingspan 7.38 m Aspect ratio 15) | 21.1 | 13.2 | 9.98 | 0.547 | 1.80 |
| *Argentavis magnificens* | 13.4 | 14.6 | 7.80 | 0.957 | 1.80 |
| *Pteranodon* | 23.0 | 16.3 | 13.6 | 0.621 | 2.20 |
| *Pteranodon* (high drag) | 12.2 | 11.8 | 13.6 | 0.946 | 2.20 |
| *Quetzalcoatlus* | 15.5 | 22.0 | 16.8 | 1.24 | 2.20 |
| *Quetzalcoatlus* (high drag) | 8.21 | 15.9 | 16.8 | 1.71 | 2.20 |
| magnificent frigatebird | 19.5 | 7.95 | 3.37 | 0.357 | 1.80 |
| California condor | 13.0 | 13.5 | 6.50 | 0.915 | 1.80 |
| brown pelican | 17.0 | 10.7 | 5.32 | 0.550 | 1.80 |
| black vulture | 13.1 | 11.8 | 5.03 | 0.790 | 1.80 |
| white strok | 16.1 | 11.3 | 5.69 | 0.617 | 1.80 |
| kori bustard | 13.0 | 16.8 | 10.1 | 1.13 | 1.80 |

**Table S5. Thermal soaring performances where $C_L^*$ is the maximum lift coefficient assuming a wing-drag minimizing wingspan.** Maximum glide ratio, and horizontal speed at maximum glide ratio limiting radius, minimum sinking speed, and lift coefficients of circling envelopes assuming a wing-drag minimizing wingspan.

| Species | Maximum glide ratio | Horizontal speed at maximum glide ratio [m/s] | Limitting radius [m] | Minimum sinnking speed [m/s] | Lift coefficient of circling envelope $C_L$* |
|---|---|---|---|---|---|
| Pelagornis sandersi (Mass 21.8 kg Wingspan 6.06 m Aspect ratio 13) | 19.6 | 11.4 | 6.96 | 0.510 | 1.80 |
| Pelagornis sandersi (Mass 21.8 kg Wingspan 6.06 m Aspect ratio 14) | 20.4 | 11.6 | 7.52 | 0.500 | 1.80 |
| Pelagornis sandersi (Mass 21.8 kg Wingspan 6.06 m Aspect ratio 15) | 21.1 | 11.8 | 8.04 | 0.492 | 1.80 |
| Pelagornis sandersi (Mass 21.8 kg Wingspan 6.13 m Aspect ratio 13) | 19.6 | 11.3 | 6.81 | 0.504 | 1.80 |
| Pelagornis sandersi (Mass 21.8 kg Wingspan 6.13 m Aspect ratio 14) | 20.4 | 11.5 | 7.35 | 0.494 | 1.80 |
| Pelagornis sandersi (Mass 21.8 kg Wingspan 6.13 m Aspect ratio 15) | 21.1 | 11.7 | 7.85 | 0.486 | 1.80 |
| Pelagornis sandersi (Mass 21.8 kg Wingspan 6.40 m Aspect ratio 13) | 19.6 | 10.8 | 6.25 | 0.482 | 1.80 |
| Pelagornis sandersi (Mass 21.8 kg Wingspan 6.40 m Aspect ratio 14) | 20.4 | 11.0 | 6.72 | 0.474 | 1.80 |
| Pelagornis sandersi (Mass 21.8 kg Wingspan 6.40 m Aspect ratio 15) | 21.1 | 11.2 | 7.21 | 0.465 | 1.80 |
| Pelagornis sandersi (Mass 21.8 kg Wingspan 7.38 m Aspect ratio 13) | 19.6 | 9.36 | 4.70 | 0.418 | 1.80 |
| Pelagornis sandersi (Mass 21.8 kg Wingspan 7.38 m Aspect ratio 14) | 20.4 | 9.54 | 5.06 | 0.411 | 1.80 |
| Pelagornis sandersi (Mass 21.8 kg Wingspan 7.38 m Aspect ratio 15) | 21.1 | 9.71 | 5.43 | 0.404 | 1.80 |
| Pelagornis sandersi (Mass 40.1 kg Wingspan 6.06 m Aspect ratio 13) | 19.6 | 15.5 | 12.8 | 0.691 | 1.80 |
| Pelagornis sandersi (Mass 40.1 kg Wingspan 6.06 m Aspect ratio 14) | 20.4 | 15.8 | 13.8 | 0.678 | 1.80 |
| Pelagornis sandersi (Mass 40.1 kg Wingspan 6.06 m Aspect ratio 15) | 21.1 | 16.0 | 14.8 | 0.667 | 1.80 |
| Pelagornis sandersi (Mass 40.1 kg Wingspan 6.13 m Aspect ratio 13) | 19.6 | 15.3 | 12.5 | 0.683 | 1.80 |
| Pelagornis sandersi (Mass 40.1 kg Wingspan 6.13 m Aspect ratio 14) | 20.4 | 15.6 | 13.5 | 0.670 | 1.80 |
| Pelagornis sandersi (Mass 40.1 kg Wingspan 6.13 m Aspect ratio 15) | 21.1 | 15.8 | 14.4 | 0.659 | 1.80 |
| Pelagornis sandersi (Mass 40.1 kg Wingspan 6.40 m Aspect ratio 13) | 19.6 | 14.6 | 11.5 | 0.654 | 1.80 |
| Pelagornis sandersi (Mass 40.1 kg Wingspan 6.40 m Aspect ratio 14) | 20.4 | 14.9 | 12.4 | 0.642 | 1.80 |
| Pelagornis sandersi (Mass 40.1 kg Wingspan 6.40 m Aspect ratio 15) | 21.1 | 15.2 | 13.3 | 0.631 | 1.80 |
| Pelagornis sandersi (Mass 40.1 kg Wingspan 7.38 m Aspect ratio 13) | 19.6 | 12.7 | 8.65 | 0.567 | 1.80 |
| Pelagornis sandersi (Mass 40.1 kg Wingspan 7.38 m Aspect ratio 14) | 20.4 | 12.9 | 9.31 | 0.557 | 1.80 |
| Pelagornis sandersi (Mass 40.1 kg Wingspan 7.38 m Aspect ratio 15) | 21.1 | 13.2 | 10.0 | 0.547 | 1.80 |
| *Argentavis magnificens* | 13.4 | 14.6 | 7.80 | 0.957 | 1.80 |
| *Pteranodon* | 23.0 | 16.3 | 13.6 | 0.621 | 2.20 |
| *Pteranodon* (high drag) | 12.2 | 11.9 | 13.6 | 0.946 | 2.20 |
| *Quetzalcoatlus* | 15.5 | 22.0 | 16.8 | 1.24 | 2.20 |
| *Quetzalcoatlus* (high drag) | 8.21 | 15.9 | 16.8 | 1.71 | 2.20 |
| magnificent frigatebird | 19.5 | 7.95 | 3.37 | 0.357 | 1.80 |
| California condor | 13.0 | 13.5 | 6.50 | 0.915 | 1.80 |
| brown pelican | 17.0 | 10.7 | 5.32 | 0.550 | 1.80 |
| black vulture | 13.1 | 11.8 | 5.03 | 0.790 | 1.80 |
| white strok | 16.1 | 11.3 | 5.69 | 0.617 | 1.80 |
| kori bustard | 13.0 | 16.8 | 10.1 | 1.13 | 1.80 |

**Table S6. Thermal soaring performances where $C_L^*$ is the maximum lift coefficient assuming a fixed wingspan.** Maximum glide ratio, and horizontal speed at maximum glide ratio limiting radius, minimum sinking speed, and lift coefficients of circling envelopes assuming a fixed wingspan.
